# A stress-NRF2 response axis polarises tumor macrophages and undermines immunotherapy

**DOI:** 10.1101/2025.07.02.662726

**Authors:** Dominik J. Schaer, Nadja Schulthess-Lutz, Livio Baselgia, Kahrisan Kunasingam, Rok Humar, Kerstin Hansen, Florence Vallelian

**Affiliations:** Department of General Internal Medicine, University Hospital and University of Zurich, Zurich, Switzerland

## Abstract

Tumor-associated macrophages (TAMs) can switch between immune-activating and cancer-promoting states; yet, the stress pathways that lock them into pro-cancerous states remain obscure. In MC38 colon tumors, repeated anti-CD40 or radiotherapy created necrosis that split TAMs into peripheral Cxcl9+ and peri-necrotic Spp1+ subsets. Spatial transcriptomics, single-cell RNA-seq, and Keap1-deficient mice showed that the latter are NRF2^high^ “stress-TAMs”, with immunosuppressive and tumor-promoting activity. The same NRF2 activation gradient separates pro-inflammatory CXCL9+ and anti-inflammatory SPP1+ TAMs across diverse human cancers. NRF2^high^ TAMs silence IFN-STAT1 programmes, lose MHC-II and chemokine expression, fail to expand T cells, drive tumor cell invasion in 3D co-cultures, and foster metastasis. Constitutive hematopoietic NRF2 activation accelerated the growth of therapy-naïve MMTV-PyMT breast tumors and markedly impaired anti-CD40 efficacy in MC38 subcutaneous and lung-metastasis models. Conversely, macrophage-specific Nrf2 deletion restored immunogenic TAMs and potentiated anti-CD40 and anti-PD-1 treatments. Thus, NRF2 constitutes a stress-response axis that fixes TAMs in a pro-cancer, therapy-resistant state; inhibiting it could revive the macrophage–T–cell amplification loop and broaden immunotherapy responses.

**GRAPHICAL ABSTRACT:** 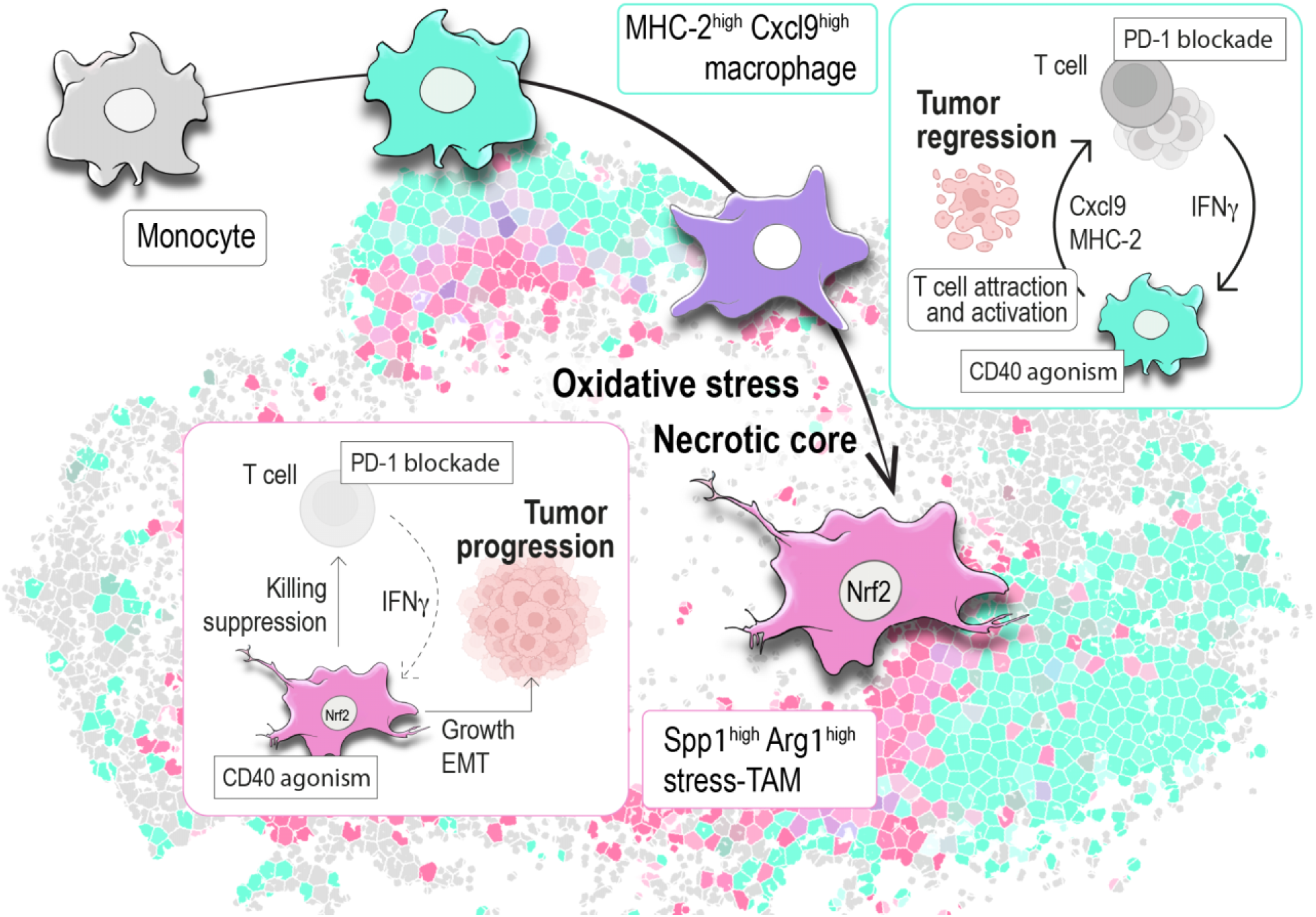

## INTRODUCTION

Tumors are heterogeneous ecosystems where diverse immune cell populations coexist and interact within distinct “niches”.^1^ Among these, tumor-associated macrophages (TAMs) are often abundant in solid tumors, showing diversified phenotypes, transcriptional programs, and functions.^2^ TAMs can either be pro-inflammatory, promoting immune-mediated tumor regression, or adopt immunosuppressive programs that foster cancer progression. These opposing polarization states clinically correlate with disease outcomes in multiple malignancies.^3^ Despite ongoing efforts to therapeutically reprogram endogenous TAMs or stem-cell derived engineered macrophages into cancer-fighting effectors ^4–7^, the specific signaling mechanisms that subvert pro-inflammatory macrophage states remain poorly characterized.^8,9^ This challenge prompts a central question: How do therapy-induced changes in the TME shape distinct macrophage subsets, and which pathways can we manipulate to suppress procancerous polarization while favoring pro-immune, anticancerous macrophages?

A growing body of evidence implicates tissue stressors—such as oxidative stress, hypoxia, acidosis, and hemorrhage—in driving TAMs toward immunosuppressive phenotypes characterized by diminished tumoricidal capacity, pro-angiogenic activity, and a propensity to facilitate epithelial-mesenchymal transition (EMT), invasion, and metastasis.^10–17^ Therapeutic interventions, such as immunotherapies and high-dose radiotherapy, exacerbate these stress conditions by inducing cell death and hemorrhage, forcing macrophages to adapt.^8,16^ A primary mediator of stress adaptation is the transcription factor NRF2 (NFE2L2).^18,19^ Under basal conditions, NRF2 is negatively regulated by KEAP1; however, upon oxidative or electrophilic stress, NRF2 stabilizes and translocates to the nucleus, driving a cytoprotective transcriptional response.^20^ While NRF2 protects macrophages against oxidative insults, it concurrently impairs their antitumor capacity, diverting them from pro-inflammatory functions to an immunosuppressive state.^21–24^ In hemorrhagic tumor areas, for example, TAMs encountering damaged red blood cells and heme upregulate NRF2-responsive genes, biasing them toward procancerous phenotypes.^10; 25–28^ Consequently, macrophage-based stress resilience through NRF2 may inadvertently fuel immunosuppressive TAM states, advancing cancer progression and undermining immunotherapies.

Here, we show that immunotherapeutic interventions expand TAM phenotypic diversity along a bifurcated polarization trajectory: immune-activated Cxcl9^+^ macrophages accumulate at the tumor periphery, whereas immunosuppressive Spp1^+^ macrophages localize to perinecrotic regions where NRF2 activity is high. Our work identifies NRF2 as a driver of therapy-resistant, cancer-promoting TAM subsets and uncovers a mechanistic link between stress adaptation and immunotherapy resistance in the TME. We further identify that modifying NRF2 could restore immunogenic macrophage subsets, thereby enhancing immunotherapy outcomes. This is the first comprehensive dissection of how immunotherapy-related necrosis drives an NRF2-centered macrophage program that subverts T-cell infiltration and tumor control.

## RESULTS

### Rebound Polarization of Monocyte-derived Tumor Macrophages Under Repeated Anti-CD40 Immunotherapy

To investigate stress-induced TAM phenotypes upon immunotherapy, we treated colon carcinoma MC38 tumor-bearing mice with a single dose of agonistic anti-CD40 or isotype control antibodies on day 11 (1× anti-CD40) or with repeated doses on days 11, 13, and 15 (3× anti-CD40). The repeated-dose protocol induces extensive necrosis with patchy zones of dense leukocyte infiltration in MC38 tumors (**Figure 1A**). Tumors were harvested twenty-four hours after the last injection. Anti-CD40 injection significantly elevated the number of tumor-infiltrating CD45^+^F4/80^+^ macrophages (**Figure 1B**). We isolated these CD45^+^F4/80^+^ TAMs by FACS for single-cell RNA sequencing (scRNA-seq) and subsequent principal component analysis (PCA) to capture the primary sources of variation across samples (**Figure 1C–D**). The sample distribution along the first principal component (PC1) was consistent with an acute immune activation induced by the single antibody injection protocol, followed by a rebound upon triple antibody injection. To identify which genes drive this separation, we examined PC1 loadings, which measure the contribution of each gene to PC1. Genes with high positive loadings (e.g., Irf1, Ccl8, Cxcl10, Cd40, Gbp5, Gbp8) characterized the initial anti-CD40 antibody effect, consistent with an IFN–STAT1–driven, immune-activated phenotype (**Figure 1E** and **F**).^29,30^ In contrast, Spp1 exhibited a strong negative loading, aligning with the rebound. Spp1 is a marker of immunoresilient and procancerous TAMs.^31^ Therefore, this rebound raised questions about stress signals in the TME driving macrophages toward an immunosuppressive trajectory. Across all three samples, **Figure 1G** highlights the striking polarity along a phenotype gradient from immune-activated Cxcl9^+^ Cxcl10^+^ MHC-II^+^ macrophages towards Spp1^+^ TAMs.

**Figure 1.**
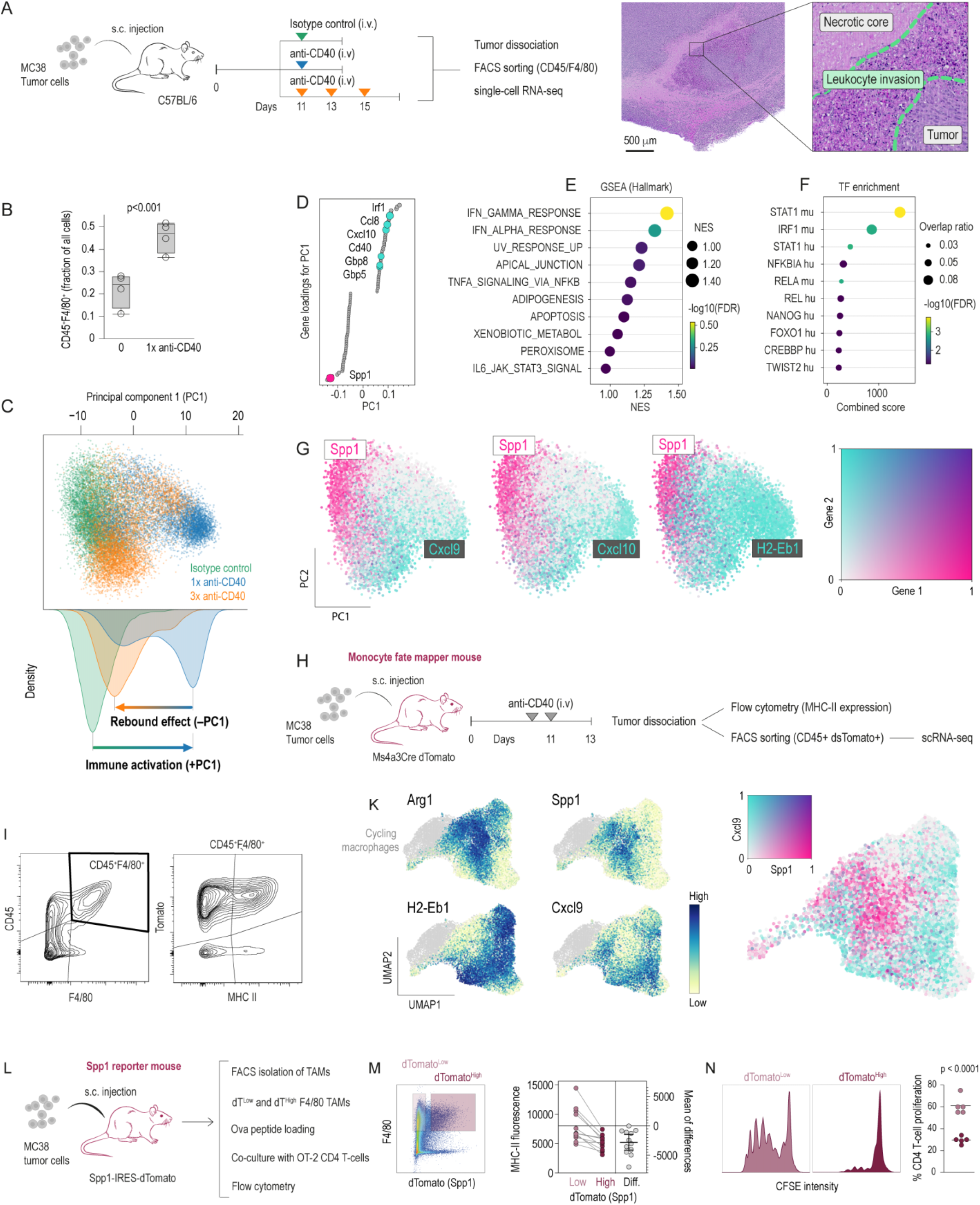
Biphasic polarization of tumor macrophages under anti-CD40 immunotherapy. **A.** Mice bearing subcutaneous (s.c.) MC38 tumors were treated intravenously (i.v.) with either isotype control or agonistic anti-CD40 antibodies on day 11 (single injection) or on days 11, 13, and 15 (multiple injections). Twenty-four hours after the final injection, tumors were harvested, and CD45^+^ F4/80^+^ cells were FACS-sorted for multiplexed scRNA-seq. Anti-CD40 therapy induces extensive tumor necrosis with patchy leukocyte infiltration as observed in the H&E-stained MC38 tumor section (right). **B.** Flow cytometry of MC38 tumor single-cell suspensions indicates increased macrophage infiltration following anti-CD40 injection. Bars represent the fraction of CD45^+^F4/80^+^ cells among total live tumor cells (n = 3 per group; mean ± SD, t-test). **C.** After demultiplexing and normalization, we selected the macrophage compartment and performed PCA on the scRNA-seq data. The scatter plot of the first two principal components (PC1, PC2) is color-coded by treatment group. Density plots for PC1 highlight that repeated anti-CD40 therapy shifts the macrophage transcriptome back toward the control-like phenotype. **D.** Gene loadings for PC1: Genes are ordered by their contribution (loading) to PC1. High positive loadings indicate genes strongly associated with the positive PC1 direction (e.g., Cxcl10), whereas negative loadings highlight genes such as Spp1. **E**. Gene set enrichment analysis (GSEA): The top PC1-loading genes (positive direction) were subjected to GSEA (Hallmark gene sets). Dots are color- and size-coded by −log_10(FDR) and normalized enrichment score (NES), respectively. **F.** Transcription factor (TF) overrepresentation: Positive PC1-loading genes were further analyzed for TF-binding motifs. The dot plot displays key TFs (e.g., STAT1) enriched among these genes. **G.** Scatter plots of PC1 vs. PC2 are color-coded for Spp1/Cxcl9, Spp1/Cxcl10, or Spp1/H2-Eb1 expression. These data illustrate a polarity between Spp1^+^ immunosuppressive and Cxcl9^+^/Cxcl10^+^ immunogenic macrophages. **H.** Monocyte origins: Mice bearing MC38 tumors were injected with anti-CD40 on day 9 and 11, followed by tumor harvest. CD45^+^dsTomato^+^ myeloid cells were FACS-sorted for scRNA-seq. The Ms4a3-Cre Rosa26-tdTomato fate-mapping system labels monocytes and their progeny irreversibly. **I.** Flow cytometry of tumor cell suspension shows that most tumor-associated macrophages (TAMs) in the MC38 model (Ms4a3-Cre fate-mapper) are CD45^+^ F4/80^+^ dsTomato^+^, with subsets differing in MHC class II levels. **K.** scRNA-seq UMAP of CD45^+^ dsTomato^+^ macrophages from Ms4a3-Cre mice treated with anti-CD40. Expression feature plots for Arg1, Spp1, H2-Eb1, and Cxcl9 confirm a bifurcated trajectory to immunosuppressive (Spp1^+^, Arg1^+^) and pro-inflammatory (Cxcl9^+^, H2-Eb1^+^) macrophages. **L.** MC38 tumors were grown in Spp1-IRES-dTomato mice and treated with anti-CD40 antibody. **M.** CD45^+^F4/80^+^ TAMs from N=12 tumors were stratified for high versus low dTomato fluorescence and MHC2 expression quantified as mean fluorescence. **N.** dTomato^high^ and dTomato^low^ CD45^+^F4/80^+^ TAMs from n=5 tumors were FACS-sorted, loaded with Ova^323–339^ peptides, and cocultured with naive CFSE-labeled CD4^+^ T-cells from OT-II mice. After 72 hours, we found less T-cell proliferation after co-culture with dTomato high versus dTomato low TAMs (paired t-test).

To confirm that both pro-inflammatory and immunosuppressive macrophage subsets arise from common monocyte precursors, we employed Ms4a3-Cre Rosa26-tdTomato fate-mapping mice (**Figure 1H**).^32^ Flow cytometry of dissociated tumors showed that most F4/80^+^ macrophages were tdTomato^+^, with subpopulations differing in MHC-II expression (**Figure 1I**). scRNA-seq of CD45^+^tdTomato^+^ monocyte progenies supported a bifurcated differentiation trajectory that yields pro-inflammatory, MHC-II–high macrophages (Cxcl9^+^, H2-Eb1^+^), as well as Spp1^+^ Arg1^+^ immunocompromised phenotypes (**Figure 1K**).

Given that repeated anti-CD40 therapy enriches for Spp1^+^ macrophages, we next employed Spp1-IRES-dTomato reporter mice to test whether Spp1 expression directly associates with reduced immunogenic capacity of TAMs (**Figure 1L**). Flow cytometric analysis confirmed that CD45^+^F4/80^+^ TAMs with high Spp1 reporter activity express lower MHC-II (**Figure 1M**) and show impaired Ova^323–339^ antigen presentation to CFSE-labeled OT-II CD4^+^ T-cells, resulting in diminished T-cell proliferation in co-culture (**Figure 1N**). Thus, Spp1^+^ macrophages in rebound-state tumors manifest a hypoimmune phenotype that correlates directly with weakened T-cell priming ability, reinforcing the role of Spp1^high^ TAMs as a marker and driver of immune evasion.

These data suggest that as therapy-induced stress intensifies, macrophages adopt an Spp1^high^ state correlated with diminished MHC-II and suboptimal T-cell priming. This “rebound” phenomenon suggested that region-dependent adaptation could counter the immune-activating signals in stressed tumor microenvironments —an inquiry we pursued by examining the spatial organization of TAMs.

### Spatial Zonation of Pro- and Anti-inflammatory Tumor Macrophages After Anti-CD40 and Radiation Therapy

To understand how local tissue factors reinforce these divergent TAM phenotypes, we investigated their spatial distribution in MC38 tumors following two anti-CD40 treatments (on days 6 and 8 post-inoculation) (**Figure 2A**). We performed high-definition spatial transcriptomics at a spatial resolution of ∼2 µm on a formalin-fixed, paraffin-embedded (FFPE) tumor section. H&E-staining of the analyzed sample revealed dense leukocyte infiltrates and a sizeable necrotic core (**Figure 2B**). After segmenting single nuclei, we identified macrophages guided by canonical markers and extracted the macrophage-specific transcriptome data for subsequent cell-by-cell analysis. Among the four macrophage clusters (**Figure 2C**), the small cluster 3 was reclassified as neutrophil granulocytes after manually inspecting the expressed genes. We classified the differentially expressed genes of the three remaining clusters as pro-inflammatory, anti-inflammatory, or neutral and calculated weighted ‘inflammation’ and ‘antigen presentation’ scores for each cluster (**Figure 2D**). Cluster 0 expressed high levels of MHC-II–associated genes (e.g., Cd74), reflecting an antigen-presenting, pro-inflammatory state, whereas Cluster 2 predominantly expressed Spp1 and Arg1, typical of anti-inflammatory macrophages. Cluster 1 exhibited moderate pro-inflammatory features, including expression of the T-cell chemoattractant Cxcl9, but lacked prominent antigen presentation.

**Figure 2.**
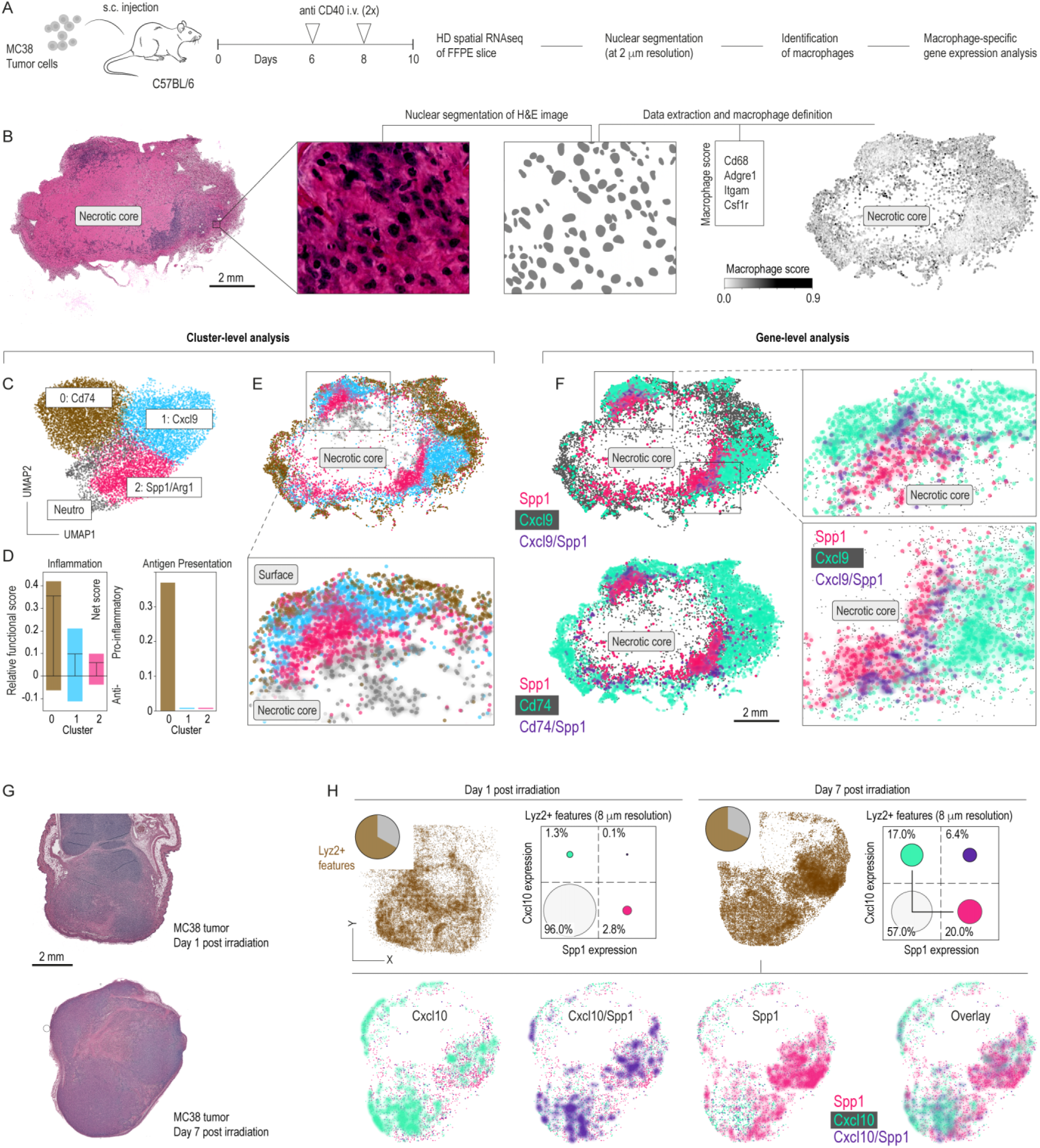
Spatial zonation of heterogeneous TAM phenotypes after anti-CD40 or radiation. **A.** Mice bearing subcutaneous (s.c.) MC38 tumors were treated intravenously (i.v.) with agonistic anti-CD40 antibodies on days 6 and 8 before they were harvested on day 10 and processed for HD spatial gene expression analysis. **B.** H&E-stained MC38 tumor section identifying a necrotic core. A higher-magnification image shows the nuclear segmentation used to achieve single-cell resolution. Macrophages were annotated using a score of marker genes. **C.** Clustering of macrophage transcriptomes reveals three subsets defined by the marker genes: Cd74, Cxcl9, and Spp1/Arg1. A small cluster (Neutro) was reclassified as neutrophil granulocytes. **D.** Weighted inflammation and antigen presentation scores per cluster indicate a pro-inflammatory (>0) versus anti-inflammatory (<0) polarization. **E.** Spatial cluster mapping of these macrophage subsets highlights distinct zonation relative to the necrotic core. Spp1/Arg1 macrophages accumulate peri-necrotically, whereas Cxcl9 and Cd74 macrophages localize toward the tumor periphery. **F.** Spatial distribution of Spp1^+^, Cxcl9^+^, and Cd74^+^ cells further emphasizes gene expression gradients within the tumor. The spatial zonation depicted here not only maps macrophage diversity but also correlates with regions likely to influence T-cell recruitment and activation. **G.** MC38-bearing mice were irradiated with 12Gy of radiation, and tumors were harvested 24 hours or 7 days post-irradiation for high-definition (HD) spatial sequencing. H&E-stained sections of these tumors are shown. **H.** Spatial RNA-seq analysis of tumor macrophages. The fractions of Lyz2^+^ features are comparable among the two samples when analyzed at 8 μm spot resolution. The comparison of day 1 and 7 post-irradiation samples shows increased phenotype polarization of macrophages along the Spp1-Cxcl10 expression axis; the size of each dot indicates the relative percentage. The 7-day post-irradiation sample was further analyzed at single-nucleus resolution, yielding spatially separated regions occupied by Spp1^+^ and Cxcl10^+^ macrophages.

The spatial mapping of the cluster identities showed a clear partition in macrophage phenotypes (**Figure 2D**). The peripheral zone was rich in macrophages classified as antigen-presenting and pro-inflammatory, while anti-inflammatory macrophages were concentrated close to the necrosis-proximal region (**Figure 2E**). In a gene-level analysis, this pattern was consistent with the accumulation of Cd74^+^ and Cxcl9^+^ macrophages at the tumor edges and Spp1^+^ macrophages clustering toward the necrotic center (**Figure 2F**). Cells co-expressing pro- and anti-inflammatory markers (Cxcl9^+^ / Cd74^+^ and Spp1^+^) were scattered between the single-positive populations, suggesting a continuous gradient of macrophage states driven by local stress.

The spatial distribution of TAMs, as revealed by our high-definition transcriptomics, not only delineates distinct regional phenotypes but also provides a framework to understand subsequent immune responses. Specifically, the accumulation of pro-inflammatory, antigen-presenting macrophages at the tumor periphery suggests a favorable niche for T-cell activation. In contrast, the concentration of Spp1^+^ macrophages near necrotic zones likely impedes effective T-cell priming and recruitment. This spatial heterogeneity sets the stage for the functional T-cell impairment presented in later sections of this paper.

Because multiple therapeutic modalities can induce necrosis, we analyzed irradiated MC38 tumors to see if they similarly exhibit Spp1^+^– Cxcl9^+^ polarization. Therefore, we conducted a spatial RNA-seq analysis on MC38 tumors 24 hours and 7 days post–irradiation with 12 Gy. **Figure 2G** shows the H&E-stained slices analyzed. As observed under anti–CD40–mediated immunotherapy, we discovered that phenotype heterogeneity among Lyz2^+^ features increased over time (**Figure 2H**), reproducing the bifurcated polarization trajectory along the Spp1–Cxcl9/Cxcl10 axis. High-resolution macrophage profiling similarly confirmed the regional segregation of Cxcl10^+^ (better detected in these samples than Cxcl9) and Spp1^+^ cells.

The convergence of rebound immunosuppression seen with repeated anti-CD40 therapy and this necrosis-proximal enrichment of Spp1^+^ TAMs in both treatments prompted us to explore how necrotic tissue damage-activated pathways, such as NRF2, might underlie macrophage plasticity in these stressed conditions.

### An NRF2-Mediated Stress Response Fuels TAM Heterogeneity

Tissue ischemia, necrosis, and injury-associated toxins such as heme are typically sensed by macrophage NRF2, triggering an adaptive response.^23, 26, 33–35^ Therefore, we hypothesized that NRF2 might orchestrate the shift toward Spp1^+^ TAMs. To characterize a prototypical phenotype of NRF2-driven TAMs, we purified macrophages from MC38 tumors grown in Keap1^flox/flox^ VavCre mice (n=4) and wild-type (WT) controls (n=5) (**Figure 3A**). In these conditional Keap1 knockout (KO) mice, NRF2 is constitutively activated in hematopoietic cells.^36^ Bulk RNA-seq of F4/80^+^ TAMs showed widespread transcriptional reprogramming consistent with persistent NRF2 activity, including elevated Gclm (a canonical NRF2 target). Importantly, the NRF2-activated TAMs reflected hallmark features of our necrosis-associated macrophages with increased Spp1 and Arg1 expression, negatively correlated with Cxcl9 and Cd74 (**Figure 3B**).

**Figure 3.**
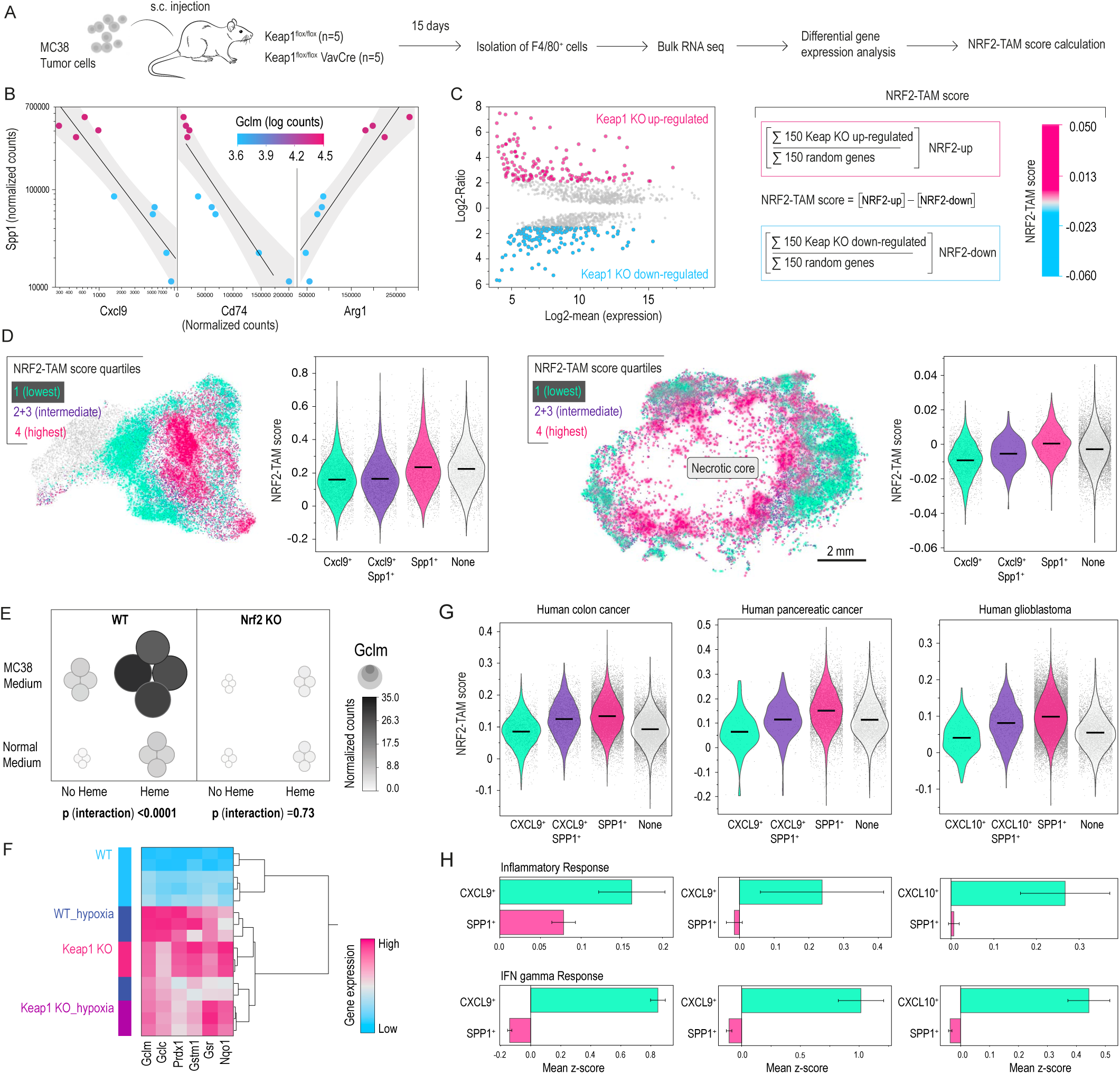
An NRF2-driven transcriptional program defines the immunotherapy-associated TAM dichotomy. **A.** MC38 tumors grown s.c. in Keap1^flox/flox^Keap1 VavCre (KO; n=4) and WT (n=5) mice were collected on day 15. TAMs (F4/80+) were purified and analyzed by bulk RNA-seq. **B.** Correlation analysis of Spp1, Cxcl9, Cd74, and Arg1 expression in Keap1 KO vs. WT TAMs, with color-coded Gclm (an NRF2 target). Keap1 loss strongly enhances Gclm expression and shifts TAMs toward an immunosuppressive phenotype (high Spp1/Arg1, low Cxcl9/Cd74). **C.** MA plot showing log-fold change vs. average expression of differentially expressed genes in Keap1 KO vs. WT TAMs. The 150 most over- and under-expressed genes define a gene signature used to calculate a score for NRF2^high^ stress-TAMs. **D.** NRF2^high^ stress-TAMs scores were calculated cell-by-cell for the scRNA-seq data in Figure 1K (left) and the spatial RNA-seq data in Figure 2 A-E (right). Features within the highest and lowest score quartiles were highlighted in the UMAP and spatial maps, respectively, mirroring the geography of macrophage polarization states. Violin plots visualize increasing NRF2^high^ stress-TAM scores in the order of Cxcl9^+^, Cxcl9^+^/Spp1^+^, and Spp1^+^ features (ANOVA with Tukey–Kramer, p<0.001 for all pair-wise comparisons). **E.** BMDMs (WT vs. Nrf2 KO) were treated with regular or MC38-conditioned medium ± heme. Regression modeling indicated strong synergistic effects (p for interaction heme x MC38 medium) of the two stimuli on Gclm expression in WT but not Nrf2 KO macrophages. Gene expressions were measured by RT-qPCR and normalized to Hprt. Each bubble or dot represents macrophages from one animal. **F.** Hierarchical clustering of canonical NRF2 target genes in Keap 1 or WT BMDMs cultured for 96 h under normoxia (21 % O₂) or hypoxia (0.2 % O₂). Each row represents an independent bulk-RNA-seq sample (3 to 5 biological replicates per condition). The heat map displays column-wise, normalised log₂-transformed counts; magenta indicates high gene expression, and blue indicates low gene expression. WT hypoxia and Keap1 KO samples group within the same cluster, suggesting that both conditions strongly enhance the expression of NRF2 target genes. **G.** Human scRNA-seq reanalysis of colon (341170 cells, 205 samples), pancreatic (12200 cells, 20 samples from two studies), and glioblastoma cancers (n=30864 cells, 31 samples) reveals that SPP1^+^ macrophages have a higher NRF2^high^-TAM score, whereas lower scores align with CXCL9^+^/CXCL10^+^/CD74^+^ subsets. (ANOVA with Tukey‒Kramer posttest, p<0.001 for all pair-wise comparisons). **H.** Inflammatory response and interferon gamma response scores (hallmark gene sets) were calculated for each cell. Bars show the mean ± 95% CI per subset within each tumor type. CXCL9⁺ and CXCL10⁺ macrophages exhibit significantly higher inflammatory and IFN-γ signatures than SPP1⁺⁺ macrophages in all three cancers.

We called these TAMs “NRF2^high^ stress-TAMs” and built a signature of NRF2-directed macrophages based on the top 150 most differentially expressed genes (up- and downregulated) between Keap1 KO and WT TAMs (**Figure 3C**). We have used this signature to classify the macrophages identified in our MC38 tumor TAM fate mapping study (**Figure 1K**) into NRF2-TAM score quantiles. The UMAP in **Figure 3D** reveals that the highest quartile scores overlap with Spp1^+^ macrophages, and across the entire data set, Spp1^+^ macrophages had higher NRF2-TAM scores than Cxcl9^+^ cells. In another analysis, we projected the cell-by-cell calculated NRF2-TAM scores onto the spatial transcriptomics dataset analyzed in Figure 2. This revealed a spatial organization with the highest NRF2-TAM scores close to the necrotic core, again, overlapping strongly with Spp1^+^ macrophages (**Figure 3D**). Also, in this case, cells expressing Spp1 matched high NRF2-TAM scores, whereas Cxcl9^+^ macrophages had lower scores.

To experimentally establish that NRF2 activation occurs in the tumor tissue-stress context, we turned to an in vitro system. First, bone marrow-derived macrophages (BMDMs) from Nrf2 KO and WT mice were exposed to MC38-conditioned medium and the necrosis-toxin heme, which synergistically induced the NRF2 target gene Gclm in WT but not Nrf2 KO BMDMs. Second, we cultured WT and Keap1 KO BMDMs for 96 hours in hypoxia (0.2% O₂). Hypoxia strongly elevated the expression of canonical NRF2-target genes in WT cells, reaching the expression observed in NRF2-activated Keap1 KO macrophages. These observations suggest that archetypal TME stresses broadly activate NRF2 in macrophages (**Figure 3F**).

To generalize our observation that the dichotomous Cxcl9^+^ and Spp1^+^ TAM spectrum reflects an NRF2 activation gradient, we reanalyzed publicly available scRNA-seq data from human colorectal and pancreatic cancers as well as glioblastoma (**Figure 3G**).^37^ Consistent with our observations in the mouse model, we identified the same pattern across these human cancer types: CXCL9^+^ macrophages with broadly pro-inflammatory and IFN gamma-activated gene expression profiles (**Figure 3H**) exhibited the lowest NRF2^high^ stress-TAM scores; anti-inflamatory SPP1^+^ macrophages displayed the highest scores, and double-positive cells showed intermediate scores.

Collectively, these findings establish a model in which microenvironmental stress drives the transition of macrophage phenotypes from pro-inflammatory to anti-inflammatory states. This macrophage phenotype shift may influence macrophage immune functions, their interactions with tumor cells, and therapeutic responses.

### NRF2 Drives Anti-Inflammatory Stress Macrophages

Next, we evaluated the function of NRF2^high^ stress-TAMs and how they respond to immunostimulatory cues by performing scRNA-seq of CD45^+^ tumor-infiltrating leukocytes from Keap1^flox/flox^ VavCre and WT mice (n=2 per group) (**Figure 4A**). As shown in **Figure 4B**, an integrated UMAP of scRNA-seq data demonstrates that macrophages, which were the most abundant leukocyte population, cluster distinctly in WT vs. Keap1 KO mice, indicating a genotype-dependent shift in their transcriptome. Differential expression, gene set enrichment analysis (GSEA), and cell-by-cell functional scoring using published gene sets for specific macrophage functions^40^ indicated a pronounced upregulation of oxidative-stress defense and metabolic adaptation in Keap1 KO macrophages. This was accompanied by reduced MHC-II expression, impaired antigen presentation capacity, and suppressed interferon-response pathways, which are central signaling pathways of anti-tumor macrophage reprogramming^7,41^ (**Figure 4C–D**). Despite significant shifts in oxidative stress defense and antigen presentation, key macrophage-defining functions such as phagocytic capacity remained comparable between WT and Keap1 KO TAMs. Transcription factor motif analysis further identified NRF2 as a top-activated factor and STAT1 as a repressed factor in Keap1 KO TAMs (**Figure 4E**).

**Figure 4.**
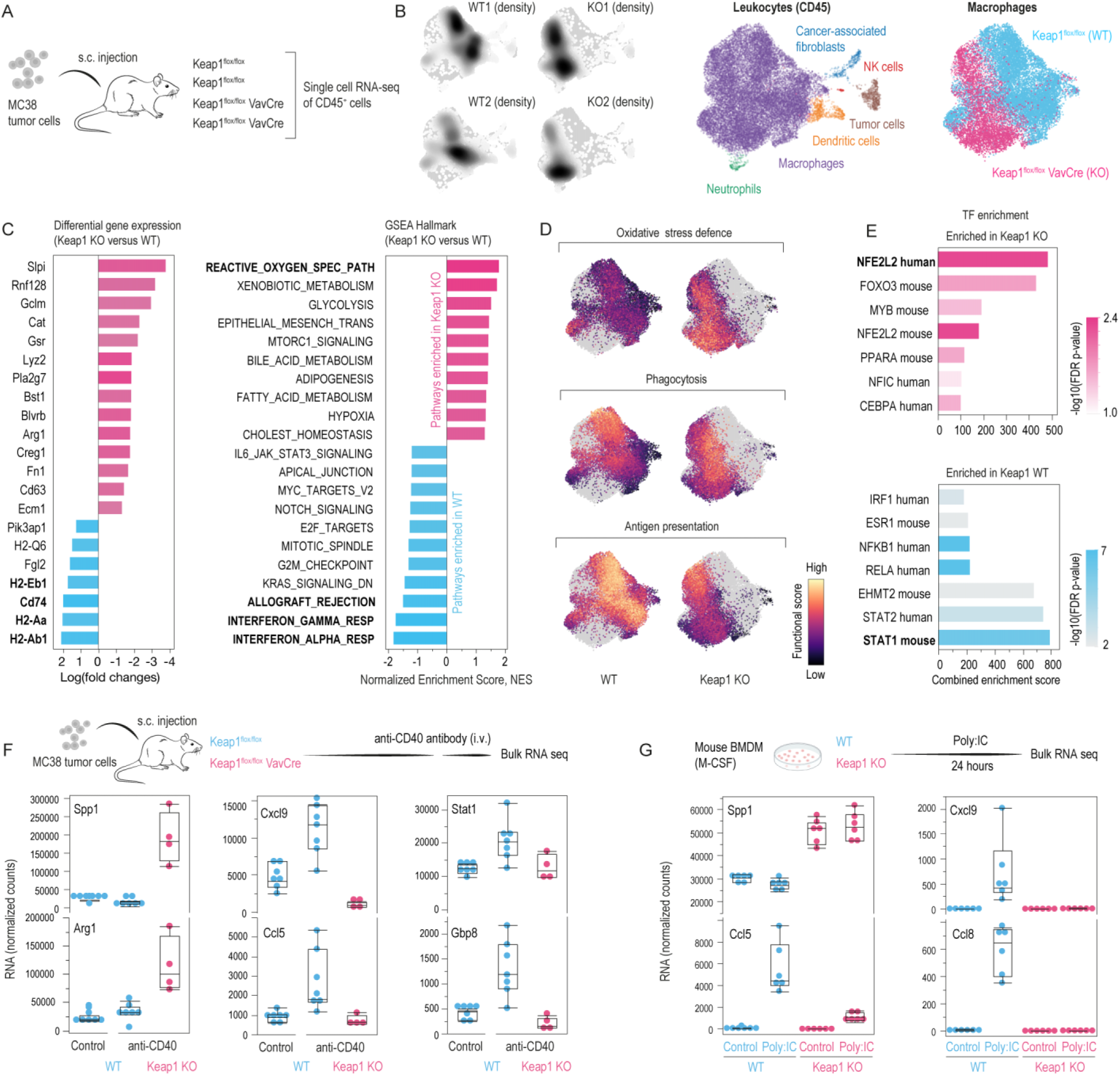
NRF2^high^ stress-TAM identity and responsiveness to immunostimulatory cues in MC38 tumors. **A.** MC38 tumors grown s.c. in Keap1^flox/flox^ VavCre (KO; n=2) or WT (n=2) mice were harvested on day 15. CD45^+^ leukocytes were analyzed by scRNA-seq to reveal NRF2-driven macrophage programs. **B.** Integrated UMAP with sample-wise cell density projections showing genotype-dependent transcriptomic shifts, with macrophages as the most abundant leukocyte subset. **C.** Differential gene expression and hallmark GSEA identify key pathways (e.g., oxidative stress defense) elevated in Keap1 KO TAMs. **D.** Functional score maps indicate increased oxidative stress defense but reduced antigen presentation in Keap1 KO vs. WT TAMs, with comparable phagocytic capacity. **E.** Transcription factor overrepresentation analysis reveals NFE2L2 (NRF2) as the top-activated and STAT1 as the repressed factor in KO TAMs, suggesting an anti-inflammatory, immunodeficient phenotype. **F.** Expression (normalized counts) of selected genes in WT vs. Keap1 KO TAMs after anti-CD40 treatment shows that NRF2 confers resistance to macrophage-activating immunotherapy. **G.** Similarly, WT vs. Keap1 KO BMDMs treated with poly(I:C) confirm that NRF2 constrains the macrophage response to inflammatory stimuli.

In an anti-CD40 therapy context, F4/80^+^ macrophages from WT tumors responded with robust induction of pro-inflammatory genes (Cxcl9, Ccl5, and interferon-responsive genes). However, Keap1 KO TAMs were largely refractory, indicating that NRF2 restricts macrophage plasticity under immunostimulatory conditions (**Figure 4F**). Parallel in vitro assays with poly(I:C)–-stimulated BMDMs confirmed the same phenomenon, demonstrating that NRF2 activation restricts macrophage responses to inflammatory stimuli (**Figure 4G**).

These findings further emphasize the role of NRF2 in driving an immune-resilient macrophage phenotype that resists reprogramming by therapeutic interventions. Given that TAM-mediated antigen presentation is crucial for T-cell priming, we next examined whether NRF2^high^ stress-TAMs influence T-cell proliferation and immunity.

### NRF2-Activated TAMs Undermine T-cell–Based Tumor Immunity

We examined the functional consequences of NRF2^high^ stress-TAM polarization for antigen-specific T-cell priming, using an antigen presentation assay with Ova^323–339–loaded^ macrophages co-cultured with CFSE-labeled OT-II (CD4^+^) T-cells (**Figure 5A**). Macrophages used in the assay were either CD45^+^ F4/80^+^ TAMs isolated from MC38 tumors or in vitro cultured BMDMs. Flow cytometry after 72 hours showed that T-cells co-cultured with WT macrophages underwent robust proliferation and upregulated the activation marker CD69. In contrast, T-cells co-cultured with Keap1 KO macrophages showed little to no CFSE dilution or CD69 induction (**Figure 5B**). We established an adoptive transfer setup to confirm this defect in vivo (**Figure 5C**). We implanted OVA-expressing MC38 cells into WT or Keap1^flox/flox^ VavCre hosts and transferred CFSE-labeled CD8^+^ T-cells from CD45.1 OT-I mice on day 7, followed by agonistic anti-CD40 antibody to boost antigen presentation. Flow cytometry analysis revealed a significant reduction in donor-derived CD45.1 CD8^+^ T-cells in the spleen and tumor-draining lymph nodes of VavCre Keap1^flox/flox^ mice, with almost no specific T-cells detected within their tumors (**Figure 5D**). Further analysis using CFSE-labeled adoptive T-cells demonstrated markedly attenuated proliferation profiles in VavCre Keap1^flox/flox^ mice compared to WT controls. The near-absence of OT-I T-cells in VavCre Keap1^flox/flox^ tumors likely reflects a combination of poor antigen presentation and diminished Cxcl9/Cxcl10-driven T-cell recruitment. This assumption is supported by the observation that across the 11’060 tumor samples in the TCGA PANCAN database (https://www.cancer.gov/tcga), CXCL9 expression predicts better CD8^+^ T-cell infiltration (**Figure 5E**), and NRF2 profoundly suppresses this chemokine.

**Figure 5.**
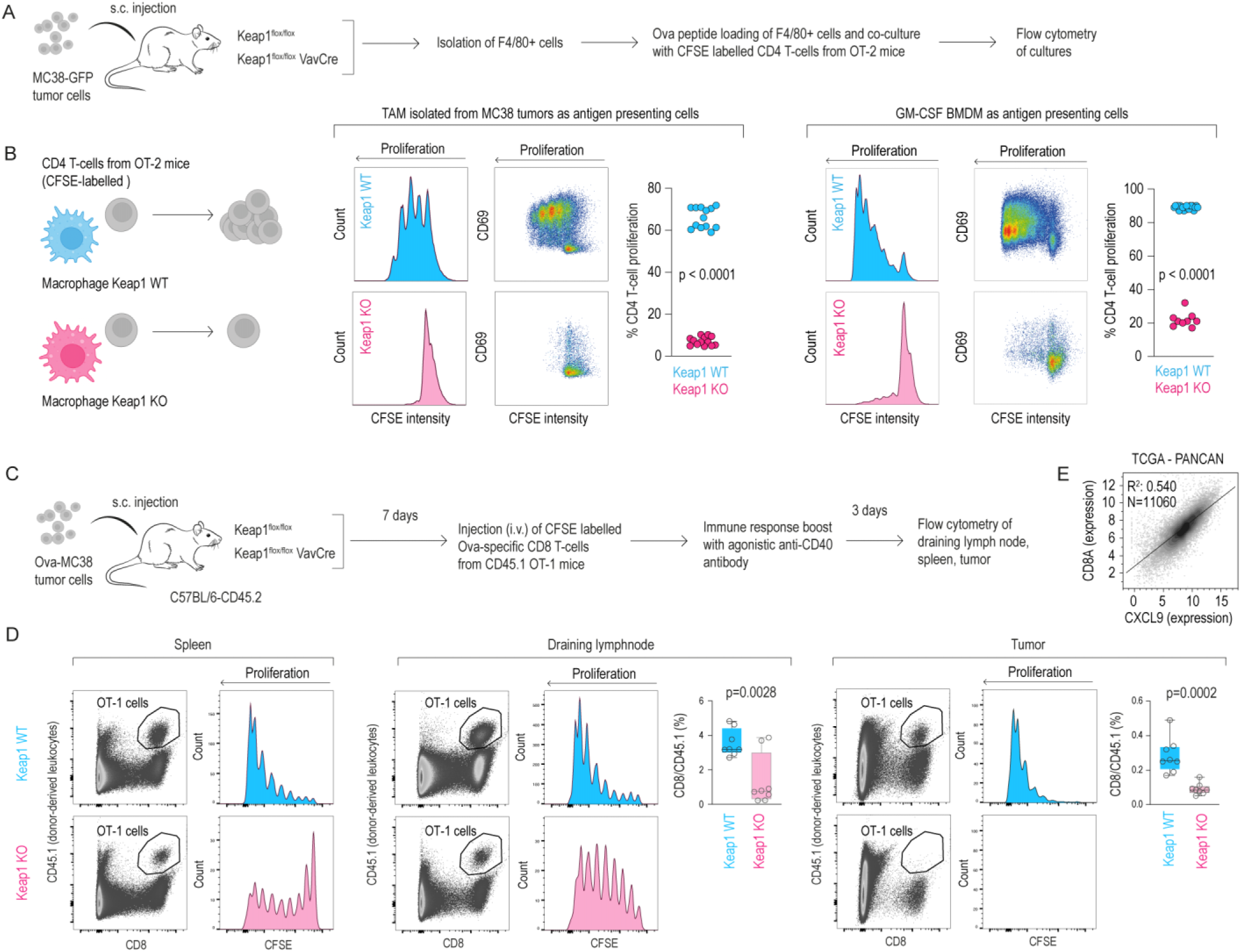
NRF2 activation in TAMs impairs antigen presentation and reduces T-cell responses. **A.** Ex vivo assay: BMDMs or MC38 tumor-derived CD45^+^F4/80^+^ TAMs from Keap1^flox/flox^ VavCre vs. WT mice were used as antigen-presenting cells after loading with Ova^323–339^ peptides. Co-cultures were performed with naive CFSE-labeled OT-II CD4^+^ T-cells. T-cell activation and proliferation (CFSE dilution, CD69 upregulation) were assessed by flow cytometry. **B.** T-cells co-cultured with WT TAMs proliferate robustly, while those with Keap1 KO TAMs show reduced CD69 expression and limited proliferation. Similar results were observed using in vitro-differentiated WT vs. Keap1 KO BMDMs **C.** In vivo assay: Ova-expressing MC38 (Ova-MC38) cells were implanted s.c. into WT or Keap1 KO mice. On day 7, OT-I CD8^+^ T-cells were injected i.v., followed by agonistic anti-CD40 antibody. T-cell proliferation and activation in the spleen, draining lymph node, and tumor were measured by flow cytometry. CD8^+^ T-cells were CFSE-labeled and genotype CD45.1 for proliferation and donor tracking, respectively. **D.** CFSE dilution profiles show reduced OT-I T-cell expansion in conditional Keap1 KO hosts. Consequently, donor-derived CD8^+^ T-cells are profoundly reduced in draining lymph nodes and primary tumors (n=7-8, mean ± SD, t-test). **E.** In the TCGA PANCAN dataset (n=11,060), high CXCL9 expression strongly correlates with increased CD8^+^ T-cell content. This suggests decreased CXCL9 expression in Keap1 KO TAMs contributes to lower CD8^+^ T-cell infiltration.

These findings indicate that the emergence of NRF2^high^ stress-TAMs severely compromises macrophage-mediated immune functions, impairing antigen presentation and restraining T-cell-mediated immunity. This macrophage NRF2-driven immune suppression highlights an overarching immunosuppressive program by which TAMs contribute to immune evasion. We next investigated whether NRF2^high^ TAMs not only impair T-cell responses but also actively reshape tumor cell behavior and the broader TME.

### Modeling Anti-inflammatory and Tumor-promoting Functions of NRF2-Driven Macrophages In Vitro

To determine whether NRF2-mediated anti-inflammatory differentiation and its downstream consequences on tumor immunology are intrinsic macrophage processes, we conducted a series of in vitro studies. First, we performed scRNA-seq studies of WT, Keap1 KO, and Nrf2 KO BMDMs after short-term culture with GM-CSF for five days to capture a broad spectrum of differentiation and polarization states. PCA analysis visualizes the significant differences in gene expression patterns between the three genotypes (**Figure 6A**, **Supplementary Figure 1**). Cell density projections along PC1 and PC2 demonstrated that Nrf2 KO cells were skewed towards MHC-II^high^ macrophages with strong antigen presentation functions. In contrast, the peak density of Keap1 KO cells was shifted away from MHC-II towards Spp1^high^ macrophages with enhanced oxidative stress functions, reinforcing the dose-dependent anti-inflammatory push provided by active NRF2 (**Figure 6B**).

**Figure 6.**
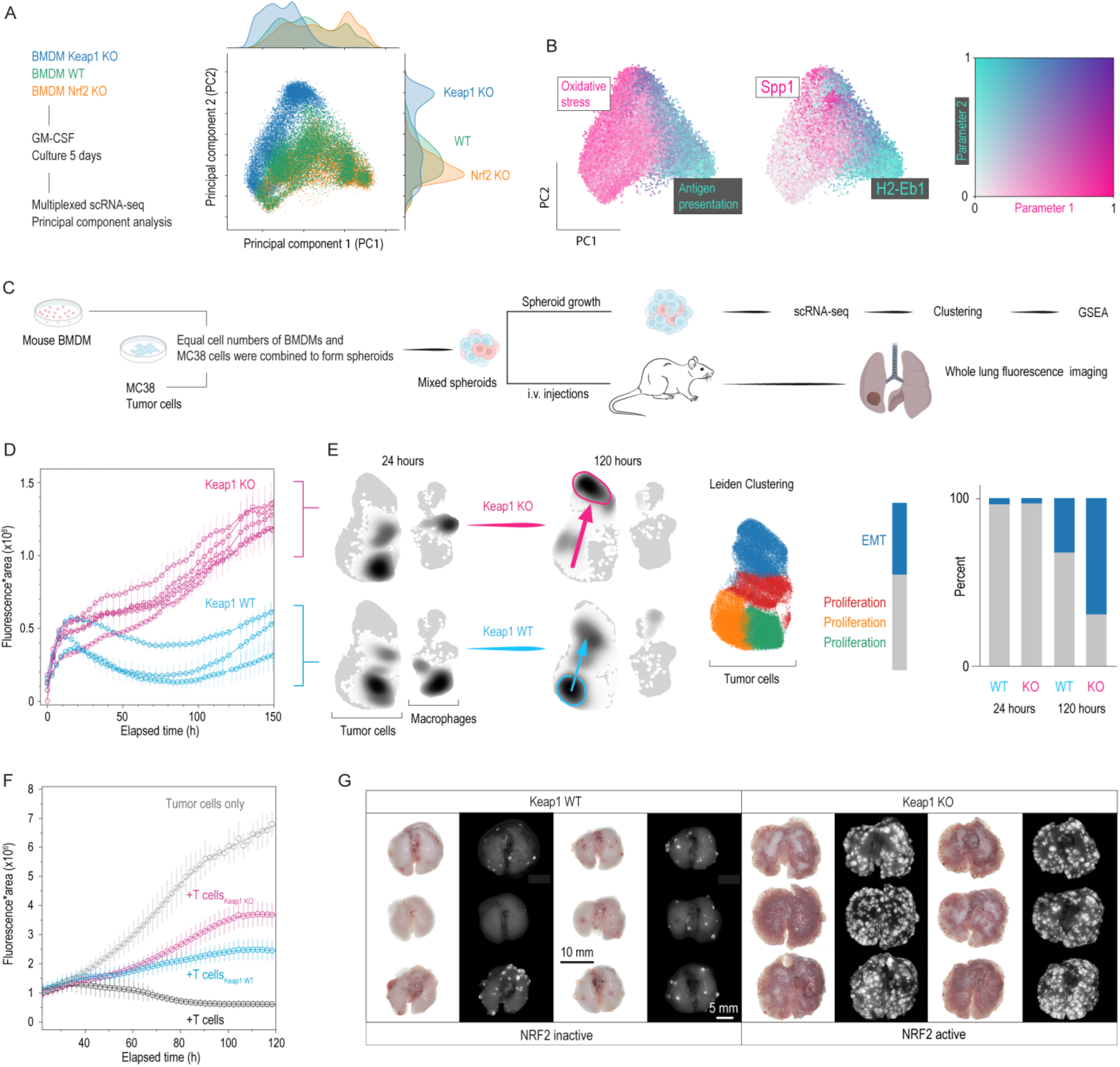
NRF2-driven TAMs promote tumor cell growth, EMT, and attenuate T-cell–mediated killing. **A.** Multiplexed scRNA-seq of GM-CSF-supported BM macrophages (WT, Nrf2 KO, Keap1 KO). The scatter plot of the first two principal components (PC1, PC2) is color-coded by treatment group. Density plots for PC1 and PC2 highlight the divergent phenotype between Nrf2 KO, Keap1 KO and WT BM cells. **B.** Scatter plots of PC1 vs. PC2 are color-coded for macrophage function scores and Spp1/H2-Eb1 expression. These data illustrate how NRF2 ablation skews BM cells from Spp1^+^ immunosuppressive towards antigen-presenting macrophages, while NRF2 activation promotes immunosuppression and loss of anti-presentation function. **C.** Spheroid co-culture model: Equal numbers of WT or Keap1 KO BMDMs were mixed with GFP-MC38 cells in ultra–low-attachment plates. Spheroid size, metastatic capacity, and T-cell killing inhibition were assessed. **D.** Live-cell microscopy tracks spheroid size (area × GFP intensity). Spheroids with Keap1 KO BMDMs grow faster than WT BMDM spheroids. Data as mean ± 95% CI (n=4 mice/genotype, 8-10 replicates each). **E.** Multiplexed scRNA-seq at 24 h and 120 h post-spheroid formation. Cell density projections on the integrated UMAP show that the cancer cell transcriptomes are very similar between genotypes despite persistent genotype-dependent variations in the macrophage transcriptome. At 120 hours, cancer cell densities undergo a marked shift, which is much more pronounced in the spheroids containing Keap1 KO macrophages. We performed Leiden clustering of the integrated data and analyzed differentially expressed genes by GSEA using the hallmark gene database. This defined that cancer cells in spheroids with WT macrophages remain mainly proliferative. In contrast, cancer cells in spheroids containing Keap1 KO macrophages progress toward an EMT state (blue cluster). The bar chart (right) quantifies the proportion of cancer cells in proliferative and EMT states. **F.** T-cell killing assay: Activated OT-I CD8^+^ T-cells were kept in regular medium or preconditioned in medium from spheroids containing either WT or Keap1 KO BMDMs, then added to GFP-OVA-MC38 monolayers. Live-cell microscopy tracked the decay of GFP fluorescence relative to the baseline growth dynamics of tumor cells. Spheroid-conditioned medium universally impairs T-cell killing, but the effect is more pronounced in Keap1 KO spheroid medium. Data represents the mean fluorescence intensity (±95% CI) of 8-10 replicates. **G.** In vivo metastasis assay: Lungs harvested three weeks after i.v. injection of ∼750 spheroids into immunodeficient mice reveal that Keap1 KO–macrophage spheroids produce more extensive metastatic disease compared to WT spheroids.

Next, we employed 3D spheroid co-cultures to explore the paracrine and direct cell-contact effects of NRF2-driven macrophages on MC38 tumor growth (**Figure 6C**). We seeded equal numbers of GFP^+^ MC38 cells and macrophages (WT or Keap1 KO) in ultra–low-adherence plates to form multicellular spheroids. Live-cell imaging showed that spheroids containing NRF2^high^ macrophages reached significantly larger sizes over 120 hours compared to spheroids with WT macrophages (**Figure 6D**). With multiplexed scRNA-seq of these spheroids collected at 24 and 120 hours post-formation, we defined the directed effects of NRF2^high^ stress-TAMs on tumor cells (**Figure 6E**). UMAP projections of the four samples visualize that the macrophage genotype did not affect tumor cell gene expression within the first 24 hours despite persistent NRF2-driven macrophage phenotypic diversity. However, by 120 hours, tumor cell transcriptomes diverged markedly, as visualized by an unequal shift in cell densities (arrows). This transcriptome shift was much more pronounced in the spheroids containing Keap1 KO macrophages. Leiden clustering and subsequent cluster-based GSEA of the tumor cell transcriptome revealed that while most tumor cells co-cultured with WT macrophages remained in a proliferative state, those co-cultured with Keap1 KO macrophages transitioned into an epithelial-mesenchymal transition (EMT) state, indicative of high metastatic potential.^42^

We further assessed whether the secretome of the macrophage-tumor spheroids may affect T-cell–mediated tumor killing (**Figure 6F**). Before measuring how activated OT-I CD8^+^ T-cells kill their OVA-MC38 target cells, we exposed them to culture medium conditioned by cancer spheroids harboring WT or Keap1 KO macrophages. MC38 proliferation and T-cell-mediated killing were recorded with an Incucyte live cell microscope. T-cells exposed to supernatants from spheroids hosting macrophages showed markedly reduced tumor cell killing, with the effect more pronounced for spheroids containing Keap1 KO macrophages. This finding indicates that NRF2^high^ stress-TAMs create an immunosuppressive microenvironment that blunts T-cell-mediated tumor lysis (**Figure 6F**).

Since EMT often correlates with metastatic potential,^42^ we next asked whether NRF2^high^ stress-TAMs influence metastasis in vivo. We intravenously injected approximately 750 macrophage-tumor cell spheroids into immunodeficient mice to assess this. After three weeks, we evaluated pulmonary metastases via GFP fluorescence imaging of the lungs (**Figure 6G**). Mice injected with spheroids containing Keap1 KO macrophages displayed significantly more extensive pulmonary metastases than those with WT macrophages.

Together, these findings demonstrate that in vitro-generated NRF2^high^ stress-TAMs adopt an anti-inflammatory phenotype, establish an immunosuppressive microenvironment, drive EMT of cancer cells, and suppress T-cell-mediated tumor cell killing, collectively contributing to metastasis and cancer progression. Having confirmed that NRF2^high^ macrophages potentiate metastasis in a reductionist setting, we next asked whether myeloid NRF2 similarly accelerates tumour progression during spontaneous, therapy-naïve carcinogenesis.

### Macrophage-intrinsic NRF2 accelerates tumour growth in a therapy-naïve setting

To determine whether leukocyte NRF2 promotes tumour progression in the absence of therapeutic stress, we crossed Keap1^flox/flox^ Vav-Cre mice with the spontaneous MMTV-PyMT breast-cancer model, a setting in which endogenous immune pressure restrains lesion outgrowth on the C57BL/6 background.^43^ Tumour latency was identical between genotypes (WT 127 ±25; KO 123.4 ±31 days), but once palpable, lesions in conditional Keap1 KO hosts expanded significantly faster than those in WT controls (Genotype × Day p = 0.0003; **Figure 7A**). Whereas 9 of 15 WT tumours entered a prolonged plateau phase, all tumors in Keap1^flox/flox^ Vav-Cre mice displayed uninterrupted growth. Thus, constitutive NRF2 activation accelerates tumor progression in a therapy-naïve context, underscoring the broad pro-tumor capacity of the NRF2–TAM axis.

**Figure 7.**
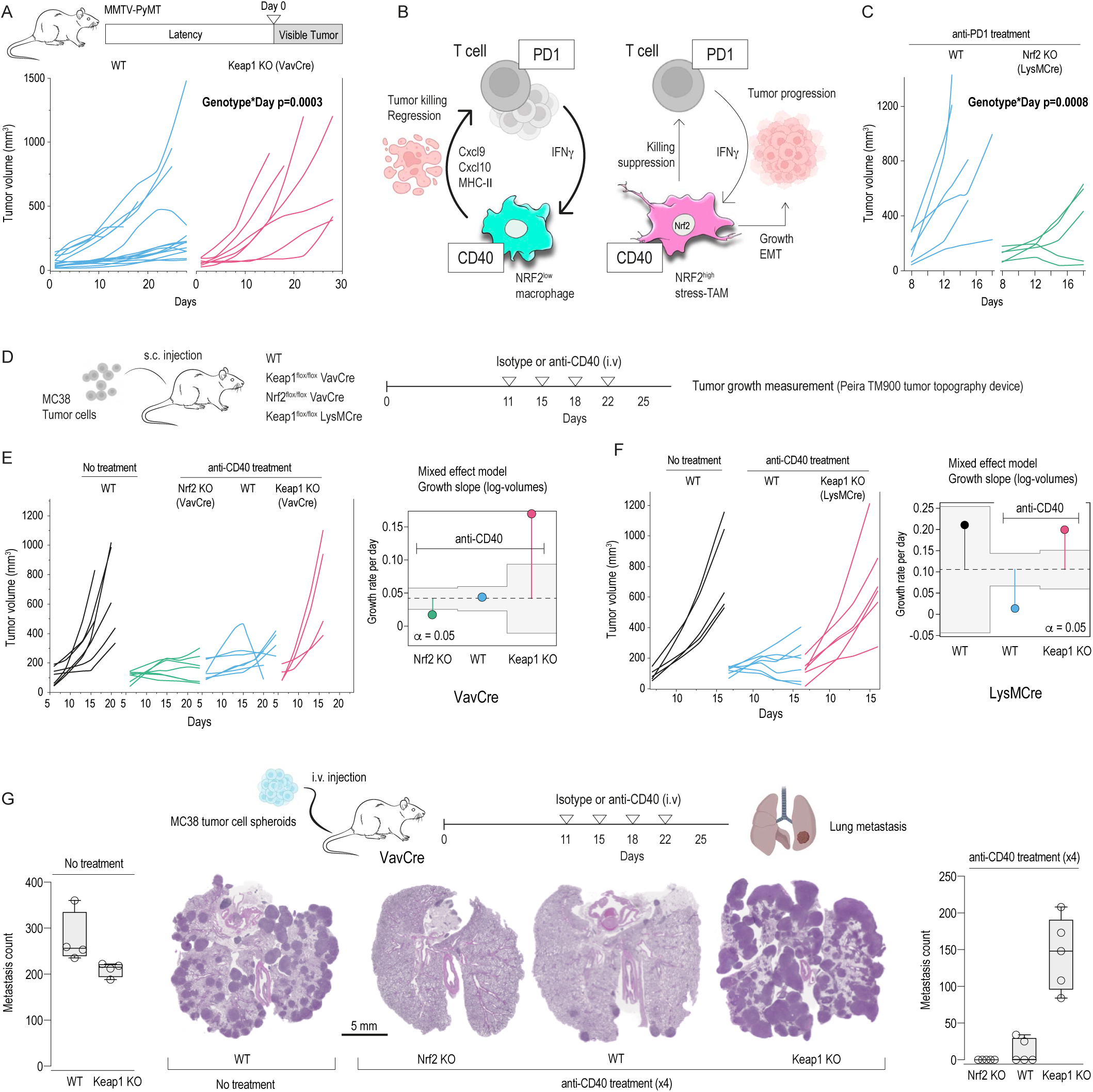
NRF2 in TAMs modulates tumor growth and the efficacy of macrophage-directed immunotherapy. **A.** Breast cancer growth was measured by digital 3D topography in WT and conditional Keap1 KO MMTV-PyMT females by an investigator blinded for genotypes. Tumors were allowed to grow until they reached 1000 mm³ or until skin necrosis occurred. The average latency to first palpable tumor was 125 days. For each animal, Day 0 indicates first tumor appearance. A mixed-effects model with Day and Genotype as fixed effects and Animal ID as random effects on log-transformed volumes reveals a significant Genotype × Day interaction (p =0.0003), indicating significantly enhanced growth rates of tumors in Keap1 KO animals. **B.** Proposed mechanistic model: Under anti-CD40 or anti-PD-1 treatments, NRF2^high^ “stress-TAMs” subvert tumor clearance, whereas reducing NRF2 activity preserves macrophage/T-cell functional interactions and enhances tumor regression. **C.** WT and macrophage-specific Nrf2-KO mice were treated with anti-PD-1 antibodies 8 days after subcutaneous MC38 inoculation. Individual tumor volumes show markedly reduced growth in Nrf2 KO. Observed vs. model-predicted log volumes confirm a good fit, and the Genotype × Day interaction (p = 0.0006) indicates significantly lower growth rates of tumors in Nrf2 KO animals. **D.** MC38 tumor-bearing mice were repeatedly treated with either isotype control or agonistic anti-CD40 antibodies i.v. Tumor volumes were measured by 3D topography. **E.** Growth curves of MC38 tumors in WT and conditional Keap1 and Nrf2 KO mice. Mixed-effect modeling of log-transformed tumor volumes with time and genotype as fixed effects and individual mice as random effects reveals significantly reduced growth slopes in Nrf2 KO and increased slopes in Keap1 KO vs. WT. UDL and LDL are upper/lower decision limits at α = 0.05. **F.** MC38 tumors in Keap1^flox/flox^ LysMCre mice also fail to respond to anti-CD40 immunotherapy, underscoring the importance of NRF2 signaling in macrophages. **G.** Pulmonary metastases were induced by i.v. injection of ∼750 MC38 spheroids and analyzed three weeks later by histology (H&E staining) and manually counting metastases. Left: Without treatment, the disease burden was similar between WT and conditional Keap1 KO mice. Right: Anti-CD40 therapy fails to control tumor spread in conditional Keap1 KO mice. Each dot represents one mouse (n=4–5 per genotype, mean ± SD, ANOVA with Tukey–Kramer post-test).

### NRF2 Targeting Enhances Immunotherapy Efficacy

Macrophages form a central nexus in a regulatory loop that can promote tumor regression through direct cytotoxicity or T cell engagement. This loop can be activated by macrophage-directed (anti-CD40) and T-cell–engaging (anti-PD-1) immunotherapies (**Figure 7B**). We conducted a series of studies to assess our hypothesis that NRF2^high^ stress-TAM phenotype shifting could undermine this immunotherapeutic paradigm.

Because PD-1 is clinically predominant, we first administered anti-PD-1 to MC38-bearing WT and Nrf2^flox/flox^ LysMCre mice on day 8 after tumor implantation. After this treatment, the macrophage-specific Nrf2-KO group displayed markedly smaller tumor volumes than WT. A mixed-effects model on log-transformed tumor volumes showed a significant Genotype × Day interaction (p = 0.0006), consistent with considerably impaired tumor growth in Nrf2-KO animals (**Figure 7C**).

Then, we switched to agonistic anti-CD40 therapy, which stimulates antigen presentation and pro-inflammatory cytokine production in macrophages.^44,45^ First, we validated the requirement of macrophages in our model by using mice lacking CD40 on macrophages (Cd40^flox/flox^ LysMCre). Compared to WT mice, these conditional KOs failed to respond effectively to anti-CD40, exhibiting larger tumors with reduced necrosis (**Supplementary Figure 2**).

Next, we monitored tumor growth in MC38-bearing mice that were repeatedly treated with anti-CD40 (**Figure 7D**). Animals included WT, Keap1^flox/flox^ VavCre (NRF2^high^), and Nrf2^flox/flox^ VavCre (NRF2-null) cohorts. Mixed-effects modeling of tumor volumes over time showed that conditional Keap1 KO mice were largely resistant to therapy, whereas conditional Nrf2 KO mice displayed improved tumor control (**Figure 7E**). Similar results were observed with MC38 tumors in Keap1^flox/flox^ LysMCre mice, underscoring that macrophage-specific NRF2 activation impairs the therapeutic effect of anti-CD40 (**Figure 7F**). Considering that the subcutaneous MC38 model may not be representative of a broader TME context, we extended these findings to a pulmonary metastasis model by injecting MC38 spheroids intravenously. Without anti-CD40 treatment, the pulmonary disease outcome after MC38 spheroid injection was not different in WT and Keap1^flox/flox^ VavCre mice, suggesting that in the absence of immunomodulatory interventions, immunosurveillance does not significantly change tumor fate within the short observation period. However, while anti-CD40 reduced lung metastasis counts in WT and nearly abolished metastases in Nrf2^flox/flox^ VavCre mice, it failed to control disease in Keap1^flox/flox^ VavCre mice (**Figure 7G**).

Collectively, our data demonstrate that ablating Nrf2 in macrophages enhances the therapeutic effect of macrophage-directed (anti-CD40) and T–cell–engaging (anti-PD-1) immunotherapies. In contrast, constitutively active NRF2 contributes to unchecked cancer progression.

## DISCUSSION

Using a combination of high-definition spatial transcriptomics, single-cell RNA sequencing, and complementary genetic models, we found that in stressed tumors, TAMs display a spatially organized spectrum of phenotypes, extending from antigen-presenting pro-inflammatory cells at the tumor periphery to anti-inflammatory macrophages enriched in necrotic regions. This distribution is consistent with a gradient of ischemia, oxidative stress, and hemorrhage, which we confirmed to be conducive to NRF2 activation. Such a gradient of macrophage states illustrates a dynamic differentiation trajectory, with monocyte-derived macrophages adapting to regional stressors as they infiltrate deeper into the tumor. By revealing that NRF2-driven macrophages cluster near necrosis and suppress T-cell immunity, our study provides new mechanistic insights into the cellular architecture and functional complexity of the TME.

Our data align with and extend prior reports that macrophage functionality is context-dependent and influenced by local metabolic, hypoxic, and oxidative conditions.^46^ After administration of anti-CD40 antibodies, which enforces pro-inflammatory macrophage reprogramming,^29,30,47,48^ monocyte-derived macrophages expressing pro-inflammatory, antigen-presentation genes accumulate at the tumor margins. In contrast, anti-inflammatory Spp1^+^/Arg1^+^ macrophages, devoid of immunostimulatory functions, cluster near necrotic cores, underscoring the significance of the “functional tumor geography.” Interestingly, a comparable macrophage phenotype dichotomy emerges under irradiated conditions, implying a shared mechanism across different clinical interventions that induce necrosis or hemorrhage. These observations underscore how therapy-related tissue damage—whether from immunotherapies or high-dose radiation—can intensify local stress, ultimately giving rise to immunosuppressive macrophage subsets.

A key mechanistic insight is the partitioning of macrophages into pro-inflammatory Cxcl9^high^/MHC-II^high^ and anti-inflammatory Spp1^high^/Cxcl9^low^/MHC-II^low^ populations along an NRF2 activation gradient. In the necrotic tumor core, where cells face persistent oxidative and metabolic insults, NRF2 stabilizes and translocates to the nucleus, enabling macrophages to survive in this hostile environment.^20^ However, while this transcriptional program is cytoprotective, our data highlight that it restricts key macrophage immune functions. Cxcl9 is an interferon-induced macrophage gene with a strong T–cell–attracting function,^49, 50^ whereas Spp1 is a potent suppressor of cytotoxic T-cells through its interactions with CD44.^51^ Across various human tumors, we found a high NRF2^high^ stress-TAM signature in Spp1+/Cxcl9- and a low NRF2^high^ stress-TAM signature in Cxcl9+/Spp1- macrophages, suggesting that NRF2-driven macrophage polarization is a universal regulatory pathway. Recent studies linking a high Spp1:Cxcl9 index to poor clinical outcomes in multiple tumor types^31^ reinforce the notion that NRF2-governed TAMs are broadly detrimental. ^52^

Tumors grown in conditional Keap1 KO mice harbor macrophages skewed towards NRF2^high^ stress-TAMs that are markedly less responsive to immune-stimulating signals. Instead of engaging in immune activation and productive cross-talk with T-cells, NRF2^high^ stress-TAMs appear locked into an anti-inflammatory, immune-suppressive state. They downregulate MHC-II expression and key interferon-responsive genes, blunting their ability to prime and recruit T-cells.(^53^ Simultaneously, NRF2^high^ stress-TAMs accelerate tumor cell proliferation, facilitate the transition of cancer cells into an EMT state, and promote a TME, which impairs T-cell killing. This synergy of tumor progression and immune evasion further illustrates how therapy-induced necrotic zones create a protected habitat for malignant cells, reflecting a barrier to effective immunotherapy.^54^ Our findings reveal that the spatial organization of TAMs is not merely a descriptive characteristic of the TME but a determinant of systemic immune responses. Pro-inflammatory macrophages at the tumor periphery foster a microenvironment conducive to T-cell priming and recruitment, whereas the immunosuppressive TAMs in necrotic cores, characterized by high NRF2 activity, correlate with reduced T-cell proliferation and function. This spatial-functional linkage underscores the importance of targeting localized stress responses to overcome systemic immunosuppression.

Critically, the tumour-promoting capacity of NRF2 extends beyond therapy-induced stress. In the spontaneous MMTV-PyMT breast-cancer model, where endogenous T-cell activity normally restrains lesion outgrowth on the C57BL/6 background,^43^ tumors in conditional Keap1 KO hosts grew rapidly once palpable, whereas >60 % of WT tumors plateaued. Latency was unchanged, indicating that NRF2 does not affect tumour initiation but rather undermines immunosurveillance during progression.

A key translational relevance of our findings lies in their implications for immunotherapy. Macrophages occupy a central nexus in a bidirectional feedback loop with T-cells: pro-inflammatory macrophages potentiate T-cell activation through enhanced antigen presentation (MHC-II) and chemokine production (Cxcl9/Cxcl10). In contrast, activated T-cells secrete IFN-γ to reinforce macrophage polarization. Thus, modulating macrophage polarization can profoundly shape T-cell fate in cancer patients—an area of immediate clinical interest.^55^ Paradoxically, treatment-related cell death, necrosis, or hemorrhage can amplify local stress, thereby activating NRF2 in macrophages.^10,27^ This negative feedback disrupts the beneficial macrophage–T-cell amplification loop: NRF2^high^ stress-TAMs dampen antigen presentation, impair T-cell infiltration and function, and eventually subvert the intended immunostimulatory effect of therapy. Indeed, our experiments showed that enforcing an NRF2^high^ stress state drastically diminished the therapeutic efficacy of anti–CD40–mediated macrophage reprogramming. In contrast, mice with macrophage-specific NRF2 deletion exhibited enhanced immunotherapeutic responses in anti-CD40- and anti–PD-1–based regimens. These observations suggest selectively targeting NRF2 or its downstream pathways in macrophages could overcome resistance and reinstate the positive T-cell–macrophage crosstalk, which is crucial for robust tumor immunity.^56^ Consistent with our anti-CD40 findings, we also found that focused radiation therapy can intensify local tissue stress, again expanding Spp1^+^ macrophages in necrotic zones, highlighting a broader phenomenon relevant to multiple treatment modalities.

We recognize several limitations of our study. First, while our mouse models and spheroid co-cultures provide mechanistic insights, the full complexity of human tumors may involve additional signals, cellular interactions, and temporal dynamics not captured here. Further in vivo experiments and longitudinal studies will be necessary to understand the timing of macrophage phenotype shifts and their reversibility. It will also be essential to validate whether interventions to modulate NRF2 activity can safely and effectively enhance immunotherapeutic outcomes in more diverse tumor models and clinical settings. Second, we identified a clear link between necrosis and NRF2-driven phenotypes. Still, we have not exhaustively delineated which local factors most robustly engage the NRF2 axis and how other transcription factors, such as hypoxia-inducible factors (HIF-1 and HIF-2), may synergize with NRF2 to further enhance metabolic reprogramming and immunosuppression.^57^ Future work must also define the downstream signaling pathways by which NRF2^high^ stress-TAMs rewire cancer cell growth, EMT state, and T-cell functions.

Our findings establish an experimental framework linking therapy-induced stress—from immunotherapy or radiation— to immune evasion via NRF2^high^ stress-TAMs. Our data reveal that the NRF2 activation state of macrophages forms a central regulatory node in a loop that can either promote tumor regression or, under NRF2-driven stress conditions, shield malignant cells from immune attack. By delineating how stress-induced macrophage states shape T-cell immunity and treatment outcomes, we provide a conceptual basis for strategies to neutralize the negative feedback imposed by NRF2^high^ stress-TAMs. Looking ahead, inhibiting NRF2 emerges as a promising tactic to restore the macrophage–T-cell amplification cycle and enhance the efficacy of immunotherapies.

## Study approval

All animal experiments were conducted under the licenses for animal experimentation approved by the Swiss Federal Veterinary Office.

## Data and code availability

Sequencing data are publicly available:

(GSE290434 [https://www.ncbi.nlm.nih.gov/geo/query/acc.cgi?acc=GSE290434]) (reviewer password: **yzqxggckdfyfzqj**)

Scripts and other materials to reproduce the data analyses can be found on Github:

https://github.com/primelab-zurich/A-stress-response-axis-in-TAMs-dictates-phenotype-heterogeneity-and-immunotherapy-evasion

Personal access token for reviewers: 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

Clone repository with token for reviewers: git clone:

https://primelab-zurich: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@github.com/primelab-zurich/A-stress-response-axis-in-TAMs-dictates-phenotype-heterogeneity-and-immunotherapy-evasion.git

## Author contributions

Conceptualization, DJS, FV; methodology, DJS, BL, FV; investigation, DJS, NSL, BL, K.K, KH, FV.; formal analysis DJS, NSL, FV; resources; visualization DJS, RH, FV; writing DJS, FV; funding acquisition DJS, FV

## Conflict-of-interest statement

The authors have declared that no conflict of interest exists.

## Acknowledgments

We thank G. Christofori and M. Spalinger at the University of Zurich for providing GFP-MC38 and OVA-MC38, respectively. Irradiated tumor samples were provided by M. Pruschy ^58^. We thank Dr. Florent Ginhoux (SingHealth and Duke NUS, Singapore) for providing the Ms4a3Cre mice.

## Funding

Swiss National Science Foundation (project grant 310030_197823 to **DJS**, 310030_201202/1 and 320030-232113 to **FV**), Swiss Cancer Foundation to **FV** (project grant KFS-5944-08-2023), Novartis Foundation to **FV**, Carigest SA. to **FV**, and Vontobel Foundation to **FV**.

## MATERIALS AND METHODS

### Animals

C57BL/6J (JAX^TM^ strain) mice were obtained from Charles River Laboratories. Ms4a3Cre mice were obtained from Dr. Florent Ginhoux (SingHealth and Duke NUS, Singapore) and bred with *Rosa26tdTomato* mice (The Jackson Laboratory). C57BL/6J-Spp1em1Msasn/J mice were obtained from Jackson Laboratories. *Conditional Keap1 knockout mice: Keap1*^tm2.Mym 36^ mice were obtained from RIKEN BRC and crossed with VavCre or LysMCre mice, which were obtained from the Swiss Immunological Mouse repository (SwImMR). *Conditional Nrf2 knockout mice:* C57BL/6-*Nfe2l2tm1.1Sred/SbisJ* (*Nrf2*^flox^) ^59,60^ mice were obtained from Jackson Laboratories and crossed with VavCre or LysMCre mice. Control littermates without the Cre driver were used for experiments involving these mouse strains. Transgenic MMTV-PyMT males in C57Bl/6 background were obtained from Jackson Laboratories and bred with VavCre Keap1 flx/flx females. Conditional Keap1 KO and WT MMTV-PyMT females were used for tumor growth experiments. *Rag2*^−/−^*γc*^−/−^ mice were obtained from the SwImMR. Other mice: B6.Cg-Tg(TcraTcrb)425Cbn/J (OT-2), C57BL/6-Tg(TcraTcrb)1100Mjb/J (OT-1), and B6.SJL-Ptprca Pepcb/BoyJ (CD45.1) were obtained from the SwImMR. CD45.1 OT-2 and CD45.1 OT-1 mice were obtained by crossing CD45.1 mice with OT-2 and OT1 mice, respectively. The CD40fl/fl mouse strain was generated from the ES clone EPD0901_3_A02, obtained from the KOMP repository (www.komp.org), by the Wellcome Trust Sanger Institute (WTSI) as described previously. ^61^. CD40 fl/fl mice were crossed with LysMCre mice.

All breeding colonies were housed and bred in the specific pathogen-free (SPF) animal facility at the Laboratory Animal Services Center (LASC) of the University of Zurich in individually ventilated cages. Male and female mice aged 7-12 weeks were used for all experiments, and all experiments with mice were performed according to animal experimentation licenses approved by the Swiss Federal Veterinary Office. For all studies, mice were randomly allocated to treatment groups, and the investigators were blinded to allocation during experiments and outcome assessment.

### Cell lines and primary cultures

#### Tumor cell line cultures

GFP-MC38 (donated by Gerhard Christofori, Department of Biomedicine, University of Basel, Basel, Switzerland) were cultured in RPMI-1640 medium supplemented with 10% fetal bovine serum (FBS, Gibco), 1% penicillin/streptomycin (P/S, Gibco), 1% nonessential amino acids (NEAA, Gibco) and 1% sodium pyruvate (Gibco). A cell line with homogeneous GFP expression was obtained by FACS sorting. Cell line authentication was performed before and after cell sorting by Short Tandem Repeat (STR) DNA genotype analysis (Microsynth, Balgach, Switzerland). GFP-OVA-MC38 (donated by Marianne Spalinger, Department of Gastroenterology, University of Zurich, Zurich, Switzerland) were cultured in DMEM medium (Gibco) supplemented with 10% fetal bovine serum (FBS, Gibco), 1% penicillin/streptomycin (P/S, Gibco), 1% nonessential amino acids (NEAA, Gibco) and 8.75ml 20% Glucose (Bichsel).

Cell cultures were maintained at 37°C and 5% CO_2_ in a humidified incubator. All cells used in this study were confirmed to be negative for mycoplasma. For all studies, mice were randomly allocated to treatment groups, and the investigators were blinded to allocation during experiments and outcome assessment.

#### BM cultures

BM cells were isolated by flushing the femurs and tibias of 7- to 12-week-old mice and then passed through a 70-μm filter. The BM cells were plated at a density of 3 × 10^5^ cells/ml on tissue culture-treated 60 mm UpCell dishes (Nunc™ UpCell™, ThermoFisher) in complete RPMI-1640 medium (10% fetal calf serum (FCS), 1% L-glutamine and 1% P/S) supplemented with 100 ng/ml recombinant mouse M-CSF (PeproTech). On day 3, half of the medium was replaced. Some cultures were treated on day 3 with 300 μM heme. Poly(I:C) (InvivoGen, 500 ng/ml) was added on day 6 for 24 hours. For experiments involving conditioned medium from tumor cells, BMDMs were seeded after washing at the end of the differentiation period in 12-well plates (TPP) in MC38 conditioned medium for 12 hours and lysed in RNA lysis buffer 1% β-mercaptoethanol for transcriptome analysis. For hypoxia experiments, BMDMs were cultured in high glucose complete RPMI-1640 medium under 0.2% O₂ concentrations from day 3 to day 7 using a workstation.

For the GM-CSF-supplemented cultures, fresh medium containing recombinant mouse GM-CSF (2×, 40 ng/ml)(PeproTech) was added on day 2. On day 3, half of the medium was removed, and new medium supplemented with GM-CSF (20 ng/ml) was added. The BM cells were harvested for analysis on day 5 from the temperature-responsive cell culture plates after cooling to room temperature. Cells were washed twice in PBS and centrifuged (300*g*, 10 min) before processing.

### Heme preparation for cell culture

Hemin (heme-chloride) was obtained from Frontier Scientific (Newark). Batches were tested endotoxin-free and prepared for cell treatments as described ^62^.

### 3D tumor spheroid production, culture, and analysis

#### Single-spheroid culture

5 × 10^3^ GFP-MC38 cells ± BMDMs (at a 1:1 ratio) were seeded in 100 μl tumor cell culture medium in 96-well Ultralow Attachment Plate PrimeSurface® 3D Culture Spheroid plates (S-BIO).

#### Multispheroid culture in microwell plates

GFP-MC38 cells (5 × 10^4^) ± BMDMs (at a 1:1 ratio) were seeded in 800 μl of tumor cell medium with M-CSF (100 ng/ml) in a 24-well SphericalPlate® 5D microwell (Axonlab). On day 3, 800 μl of fresh culture cell medium was added.

#### Quantification of spheroid growth and invasion

Single spheroids were imaged in the cell culture incubator with an IncuCyte S3 instrument (Sartorius). Green fluorescence and phase contrast images of the spheroids were acquired every 3-4 hours for seven to ten days. The area and fluorescence intensities of the images were measured using the IncuCyte Spheroid Software Module (Sartorius). Data are reported as spheroid fluorescence intensity integrated across the spheroid area (for tumor cells expressing a fluorescent protein) or as spheroid area.

### T cell assays

#### CD4+ T and CD8+ T cell isolation and CFSE labeling

Lymphocyte T cells were positively enriched from spleen single-cell suspensions using CD4 or CD8 enrichment kit (Thermofisher) according to the manufacturer’s instructions. Isolated CD4+ T or CD8+ T cells were labeled with CFSE (Thermo Fisher) at RT for 20 min, washed with RPMI Medium and counted before use. The final purity, confirmed by flow cytometry, was >95%.

#### OT-2 assay

A total of 2×10^4^ FACS-sorted CD45+ F4/80+ TAMs or GM-CSF stimulated BM cells were plated in 96-well round-bottom plates (Falcon), pulsed or not with 1 μg/ml Ova^323–339^ peptides (Sigma) for 45 min at 37 °C, and washed three times with PBS. Subsequently, 1 × 10^5^ CFSE-labeled naive CD4+ T cells isolated from spleens of OT-2 mice were added to the BM cells in complete RPMI-1640 medium and cocultured at 37 °C. CFSE dilution was assessed by flow cytometry after three days.

#### Adoptive Transfer of OT-1 CD8+ T cells

2 × 10^6^ CFSE-labeled CD8+ T cells isolated from OT-1 × CD45.1 mouse spleens were injected intravenously via the tail vein into conditional Keap1 KO mice and WT littermates. The mice were challenged intravenously with agonistic anti-CD40 antibody (20 mg/kg, InVivoPlus, clone FGK4.5). The mice were sacrificed three days later, and CD45.1+ CD8+ T cells were analyzed by flow cytometry.

#### CD8+ T cell activation

1 × 10^6^ CFSE-labeled CD8+ T cells isolated from OT-1 mouse spleens were activated with anti-CD3/anti-CD28 Dynabeads (Invitrogen) and incubated overnight in complete RPMI-1640 medium or in supernatant of mixed cell spheroids that have been cultured in microwell plates for 5 days.

#### T-cell killing assay

5 × 10^3^ GFP-OVA-MC38 cells were seeded into 96-well flat bottom plates (TTP) and cultured for 20 hours. Subsequently, 5 × 10^3^ activated OVA-specific CD8+ T cells were added to the tumor cells, and green fluorescence and phase-contrast images were acquired every 2 hours. The area and fluorescence intensities of the images were measured using the IncuCyte Cell-by-Cell Software Module (Sartorius). Data are reported as fluorescence intensity integrated across the area.

### Tissue/cell preparation and digestion

#### Tumor digestion

Subcutaneous tumors were excised and minced using a scalpel on a sterile Petri dish. The minced tumor tissue was then dissociated in 3 ml of digestion medium (RPMI medium (Gibco) supplemented with 25 µg/ml Liberase™ (Roche) and 40 µg/ml DNase I (Roche; 2000 U/ml). This mixture was incubated for 30-45 minutes in a water bath at 37°C. The dissociated tumor cells were filtered through a 70-µm cell sieve and washed with PBS containing 2 mM EDTA to halt the digestion process. Following the lysis of red blood cells using RBC lysis buffer (BioLegend), the cells were washed again and immediately utilized.

#### Macrophage isolation from digested tumors

Anti-rat IgG Dynabeads (Invitrogen) were washed and incubated with rat anti-mouse F4/80 IgG2a antibodies (BD Biosciences) and CD11b IgG2b antibodies (BioLegend) at a ratio of 3.33 µg of antibody per 50 µl of Dynabeads. Single-cell suspensions from tumors were incubated with anti-F4/80-coated (for RT-qPCR or RNA bulk sequencing experiments) or anti-CD45 coated Dynabeads (for scRNA-seq experiment) on a rotating wheel at 4°C for 30 min. After incubation, a positive selection of Dynabead-bound single-cell suspensions was performed on a DynaMag magnet (Invitrogen) with three washing steps, as the manufacturer’s instructions suggested.

#### Spheroid digestion

Spheroids were dissociated in 2 ml digestion medium (RPMI medium (Gibco) + 25 µg/ml Liberase™ (Roche) + 40 µg/ml DNase I (Roche; 2000 U/ml) and incubated for 30-45 min in a water bath at 37°C with gentle shaking every 5 min. Then, 4 ml PBS + 0.04% BSA was added to stop the digestion. Digested spheroids were used immediately.

### Mouse models

#### Lung metastasis model in mice

Approximately 750 spheroids were collected from microwell plates (equal to the content of one macro well) 4 days post-spheroid formation and injected intravenously into the tail vein of recipient mice. Agonistic anti-CD40 treatment (20 mg/kg, InVivoPlus, clone FGK4.5) or an isotype control antibody was administered intravenously according to the treatment protocol. Two or three weeks post-injection, the lungs of anesthetized mice were perfused with PBS through the right ventricle and the trachea and collected for whole-organ fluorescence imaging with a Zeiss Discovery V8 stereomicroscope and histology.

#### Tumor growth model in mice

Once confluent, GFP-MC38 tumor cells were harvested using 5 mM EDTA (Gibco) (4 min at 37°C). MC38 cells (2 × 10^6^) in culture medium were mixed with Geltrex (Thermo Fisher) and injected subcutaneously into the mouse flanks. Agonistic anti-CD40 treatment (20 mg/kg, Bio X Cell, clone FGK4.5), antagonistic anti-PD-1 (CD279) (10 mg/kg, Bio X Cell) or an isotype control antibody was administered intravenously according to treatment protocol. Mice were euthanized, and tumors were collected at defined time points after antibody administration. Tumors were then digested or fixed in 10% formalin and stored at room temperature.

Mammary tumor growth was measured by digital 3D topography in WT and conditional Keap1 KO MMTV-PyMT females. Mammary tumors were allowed to grow until they reached 1000 mm³ or until skin necrosis occurred.

#### Tumor volume measurements

All tumor volumes were measured non-invasively by an investigator blinded to treatment or genotype with a Peira TM900 device, which extracts volumes from high resolution stereo-3D images.

### Flow cytometry

Cells were preincubated with Mouse BD Fc Block™ (≤ 1 μg/million cells in 100 μl, BD Biosciences) at 4°C for 10 min. The following antibodies were purchased from BD Biosciences: anti-CD45 (clone 30-F11), anti-F4/80 (clone T45-2342), and anti-I-A/I-E (clone M5/114.15.2). The following antibodies were purchased from BioLegend: anti-CD45.1 (clone A20), anti-CD45.2 (clone 104), anti-CD4 (clone GK1.5), anti-CD8 (clone 53-6.7), and anti-CD69 (clone H1.2F3). Corresponding isotype-matched irrelevant specificity controls were purchased from BD, and BioLegend. Multiparameter analysis was performed with an LSRFortessa analyzer (BD Biosciences). The data were analyzed using FlowJo software (version 10.7.1). Cell sorting was performed on a BD FACSAria III 4L.

### Histology

#### Organ fixation for paraffin embedding and microtome sectioning

Mice were anesthetized by intraperitoneal injection of ketamine (80 mg/kg), xylazine (16 mg/kg), and acepromazine (3 mg/kg) and transcardially perfused with cold PBS. Organs were placed in 10% formalin and transferred to 70% ethanol after 24 hours before embedding in paraffin blocks. Microtome sections (2-2.5 µm) of each organ were cut for H&E staining and immunohistochemistry.

#### Immunohistochemistry

*GFP staining:* Tissue sections were incubated overnight with a goat anti-GFP antibody (Abcam) diluted 1:1000, followed by a biotinylated horse anti-goat secondary antibody (Vector) diluted 1:500. *Anti-F4/80 staining*: Tumor sections were incubated overnight with a rat anti-mouse F4/80 antibody (Bio-Rad, MCA497G) diluted 1:80, followed by a biotinylated goat anti-rat secondary antibody (Vector, BA9401) diluted 1:500.

All immunohistochemical sections were rinsed in 0.1 M phosphate buffer, pH 7.4, and incubated with diaminobenzidine (DAB, Abcam) for 2-5 min. After incubation, sections were washed in deionized water and lightly counterstained with hematoxylin solution, according to Mayer (Sigma).

#### Microscopy image acquisition and analysis

A Zeiss Axio Scan Z1 Slidescanner microscope imaged whole-lung sections and subcutaneous tumors. Images were analyzed using Qupath and ImageJ. Brightness, contrast, and color tone (for single-channel fluorescence images) were adjusted with Adobe Lightroom software version 8.1. using identical settings for all images of an experiment.

### Sequencing-based workflows and data analysis

#### Bulk RNA sequencing

RNA was extracted from macrophages using the RNeasy Mini kit (Qiagen) according to the manufacturer’s protocol, including on-column DNase I treatment. RNA quality was validated on an Agilent Technologies 4150 Tapestation using RNA Screentapes, and only samples with an RNA integrity number (RIN) of > 9 were used for sequencing. cDNA libraries were generated at the Functional Genomics Center Zurich (FGCZ) from RNA samples using the Illumina Stranded mRNA Prep ligation kit following the manufacturer’s instructions. The quality and concentration of the libraries were determined using an Agilent Technologies 4200 Tapestation with DNA Screentapes. The libraries were pooled in equimolar amounts and sequenced on an Illumina NovaSeq X Plus sequencer (paired-end 150 bp) with a depth of at least 20 million reads per sample.

#### Bulk RNA sequencing data analysis

Reads were aligned to the reference genome Ensembl GRCm38.p5 Release 91 using STAR (v2.7.0e).^63^ The quality of alignment was evaluated using Samtools (v1.9). ^64^ Counts were obtained using the featureCounts function of the Rsubread package (v1.22.2).^65^ Differential expression analysis was performed with the DESeq2 R package (v1.26.0)^66^.

#### scRNA-seq sample preparation

Single-cell suspensions were fixed following the demonstrated protocol Fixation of cells & Nuclei for Chromium Fixed RNA Profiling (10X Genomics, CG000478) and processed for long-term storage at −80°C. After storage, up to 2 Mio cells per sample were hybridized with unique single Mouse WTA probes using BC001-004 according to the demonstrated protocol Chromium Fixed RNA Profiling Reagent Kits for Multiplexed Samples (10X Genomics, CG000527). Following gene expression library construction, ready-made libraries were sequenced at the Functional Genomics Center Zurich (FGCZ) on an Illumina NovaSeq X Plus system following the recommendations of 10X Genomics.

#### scRNA-seq sample analysis

Downstream analysis was performed in Python (version 3.10.10) with Scanpy (1.9.2).

##### Read alignment

Reads were aligned to the mouse reference genome GENCODE GRCm39 (Release_M31-2023-01-30) using CellRanger (version 7.2.0).

##### Quality Control and Preprocessing

To assess the quality of the cells, the following covariates were considered: number of genes expressed in a cell (*n_genes_by_counts*), number of counts per cell (*total_counts*), and percentage of mitochondrial RNA (*pct_counts_mt*). Cells that expressed fewer than *min_genes* or more than *max_genes* were filtered out. Cells with a percentage of mitochondrial RNA greater than *max_pct_mt* were considered dead and removed from the analysis. Genes that were expressed by fewer than *min_cells* cells were excluded. See below for the cutoff values used in each experiment. The count data were normalized so that every cell has the same total count after normalization (sc.pp.normalize_total) and log(x+1) (`sc.pp.log1p`) transformed, yielding normalized expression values. For Fig.5 an algorithm based on deconvolving size factors from cell pools implemented in the R package scran (`calculateSumFactors`)^67^ was used due to the higher complexity of the experiment data.

##### Data integration

Multiplexed samples were merged into one dataset by simple concatenation. Additionally, samples from different experiments were integrated using the harmony (`sc.external.pp.harmony_integrate`) ^68^ or the scanorama algorithm ^69^, depending on the experimental setup and its complexity.

##### Dimensionality reduction and clustering

For dimension reduction, the following steps were performed using the Python package Scanpy: identifying highly variable genes (`sc.pp.highly_variable_genes`), performing PCA using highly variable genes (`sc.tl.pca`), computing the neighborhood graph (`sc.pp.neighbors`) and computing the UMAP (`sc.tl.umap`). The cells were clustered using Leiden clustering (`sc.tl.leiden`), which depends on the neighborhood graph. The resolution of the Leiden clustering was chosen so that a biologically meaningful number of clusters was produced.

##### Cell type annotation and functional classification

To identify cell types, we analyzed the expression of marker genes and other differentially expressed genes (`sc.tl.rank_genes_groups` with `method = ‘wilcoxon’`). GSEA was performed to assess functional and biological process-related differences between clusters or conditions. First, genes were ranked using the output of the Wilcoxon rank-sum test (`rank = -log10(adj. p value)*sign(logfoldchange)`) and then fed to the GSEA algorithm implemented in the Python package decoupler ^70 71^ (`decoupler.get_gsea_df`), resulting in a normalized enrichment score (NES) and a false discovery rate (FDR) per gene set. For transcription factor analysis with the TRRUST_Transcription_Factors_2019 gene set database and for scoring enrichment of gene sets per cell, the AUCell algorithm (`decoupler.run_aucell`) was used.

#### Visium CytAssist Spatial Gene Expression for FFPE

Subcutaneous mouse tumors were processed as described in the section titled organ fixation for paraffin embedding and microtome sectioning. Microtome sections on regular glass slides were deparaffinized and stained for H&E as described in Visium CytAssist Spatial Gene Expression for FFPE (10X Genomics, CG000520). Regions of interest on the tumor sections were chosen using H&E staining and aligned in the tissue slide cassette 6.5 mm. After destaining and decrosslinking, sections were immediately subjected to probe hybridization overnight followed by probe ligation, release and extension and spatial library construction as described in Visium CytAssist Spatial Gene Expression for FFPE (10X Genomics, CG000495). Ready-made libraries were sequenced at the Functional Genomics Center Zurich (FGCZ) on an Illumina NovaSeq 6000 system.

#### Visium HD CytAssist Spatial Gene Expression for FFPE

Tissues were processed as described in the section titled organ fixation for paraffin embedding and microtome sectioning. Microtome sections on regular glass slides were deparaffinized and stained for H&E as described in Visium HD FFPE Tissue Preparation Handbook (10X Genomics, CG000684). Regions of interest on the tissue sections were chosen using H&E staining and aligned in the tissue slide cassette (6.5 mm). After destaining and decrosslinking, sections were immediately subjected to probe hybridization overnight followed by probe ligation, release and extension and spatial library construction as described in the user guide Visium HD Spatial Gene Expression Reagents Kits (10X Genomics, CG000685). Ready-made libraries were sequenced at the Functional Genomics Center Zurich (FGCZ) on an Illumina NovaSeq X Plus system using paired-end 150 bp read configuration.

#### Spatial transcriptomics analysis

Downstream analysis was performed in Python (version 3.10.10) with Scanpy (1.9.2) ^72^. Each sample was processed individually.

##### Read alignment

Reads were aligned to the mouse reference genome Ensembl GRCm39 (Release_106-2022-07-05) using SpaceRanger (version 2.1.0).

##### Nuclei Segmentation and Custom Binning of Visium HD Gene Expression Data

We reassigned the barcodes to cell nuclei to extract the macrophages from the spot-based Visium HD data to create single-cell-like data. To achieve this, the cell nuclei were segmented based on a microscopy image of the H&E stained tissue as described in [https://www.10xgenomics.com/analysis-guides/segmentation-visium-hd]. After normalization (*min_percentile*, *max_percentile*) of the H&E image, a pre-trained model (StarDist2D: 2D_versatile_he) predicts the cell nuclei. Each barcode is then assigned to a cell nucleus, and the counts are aggregated. This results in a single-nucleus resolution of the data.

##### Quality Control, Preprocessing, Dimensionality Reduction

Cell nuclei that contain less than *min_total_counts* or their area is bigger than *max_area* are removed from the dataset. The data was normalized (sc.pp.normalize_total) and log(x+1)-transformed (sc.pp.log1p) to generate gene expression data.

Macrophages were annotated by scoring (sc.tl.score_genes) the gene expression of tissue-specific *macrophage marker genes*.

For further analysis, only cells labeled as macrophages were considered.

Based on the top 2000 highly variable genes (sc.pp.highly_variable_genes), a PCA was performed (sc.tl.pca), and a neighborhood graph was constructed (sc.tl.neighbors).

##### TAM-NRF2 scoring

Using the gene sets obtained from the differential gene expression analysis of bulk RNA-seq of F4/80^+^ TAMs between Keap1^flox/flox^ VavCre and WT mice, a combined TAM-Nrf2-on/TAM-Nrf2-off score was calculated (sc.tl.score_genes).

### Statistics

Data plotting and statistical analysis were performed with Prism 11 (GraphPad) and JMP 17 PRO (SAS). We used ANOVA with Tukey‒Kramer posttest to account for multiple comparisons and t-tests (two-tailed), as indicated. Tumor growth curves were analyzed with the JMP model builder using a mixed effect model, setting time, genotype, and their interaction as fixed effects and individual mice as random effects. All data points are displayed in the plots to visualize the data distribution. *P*-values are indicated in figures or stated in the text.

## Supplementary Figures

**Supplementary Figure 1**

Multiplexed scRNA-seq at 24 h and 120 h post-spheroid formation (Figure 6E). Leiden clustering of the integrated data and GSEA of the differentially expressed genes.

**Supplementary Figure 2**

**A.** MC38 tumors were grown s.c. in CD40^flox/flox^ LysM versus WT mice. On day 15, mice received anti-CD40 treatment administered i.v. Three days after the second dose, tumors were collected for in situ GFP fluorescence imaging and histology.

**B.** Left and right: Bright-field and GFP fluorescence images visualizing MC38 tumors in situ. Middle: GFP fluorescence intensity integrated across the tumor area. Each dot represents one tumor grown on the right and left flanks of a mouse (n = 12 tumors; mean ± SD, t-test).

**C.** Representative H&E-, anti-GFP and anti-F4/80 stained-MC38 tumor sections.

## Notes

### Competing Interest Statement

The authors have declared no competing interest.

## REFERENCES

1. Klein, L. et al. Spatial tumor immune heterogeneity facilitates subtype co-existence and therapy response in pancreatic cancer. Nat Commun 16, 335 (2025).

2. Pollard, J. W. Tumour-educated macrophages promote tumour progression and metastasis. Nat Rev Cancer 4, 71–78 (2004).

3. Pittet, M. J., Michielin, O. & Migliorini, D. Clinical relevance of tumour-associated macrophages. Nat Rev Clin Oncol 19, 402–421 (2022).

4. Molgora, M. & Colonna, M. Turning enemies into allies-reprogramming tumor-associated macrophages for cancer therapy. Med 2, 666–681 (2021).

5. Engblom, C., Pfirschke, C. & Pittet, M. J. The role of myeloid cells in cancer therapies. Nat Rev Cancer 16, 447–462 (2016).

6. Lei, A. et al. A second-generation M1-polarized CAR macrophage with antitumor efficacy. Nat Immunol 25, 102–116 (2024).

7. Pucci, F. & Coussens, L. M. Redirecting tumor macrophage activity to fight cancer: Make room for the next era of anti-cancer drugs. Cancer Cell 39, 1300–1302 (2021).

8. De Palma, M. & Lewis, C. E. Macrophage regulation of tumor responses to anticancer therapies. Cancer Cell 23, 277–286 (2013).

9. Tzetzo, S. L. & Abrams, S. I. Redirecting macrophage function to sustain their ‘defender’ antitumor activity. Cancer Cell 39, 734–737 (2021).

10. Schaer, D. J. et al. Hemorrhage-activated NRF2 in tumor-associated macrophages drives cancer growth, invasion, and immunotherapy resistance. J Clin Invest 134, (2023).

11. Wu, J.-Y. et al. Cancer-Derived Succinate Promotes Macrophage Polarization and Cancer Metastasis via Succinate Receptor. Mol Cell 77, 213–227.e5 (2020).

12. Chen, S. et al. Tumor-associated macrophages are shaped by intratumoral high potassium via Kir2.1. Cell Metab 34, 1843–1859.e11 (2022).

13. Devalaraja, S. et al. Tumor-Derived Retinoic Acid Regulates Intratumoral Monocyte Differentiation to Promote Immune Suppression. Cell 180, 1098–1114.e16 (2020).

14. Bohn, T. et al. Tumor immunoevasion via acidosis-dependent induction of regulatory tumor-associated macrophages. Nat Immunol 19, 1319–1329 (2018).

15. Henze, A.-T. & Mazzone, M. The impact of hypoxia on tumor-associated macrophages. J Clin Invest 126, 3672–3679 (2016).

16. Colegio, O. R. et al. Functional polarization of tumour-associated macrophages by tumour-derived lactic acid. Nature 513, 559–563 (2014).

17. Folkert, I. W. et al. An iron-rich subset of macrophages promotes tumor growth through a Bach1-Ednrb axis. J Exp Med 221, (2024).

18. Itoh, K. et al. An Nrf2/small Maf heterodimer mediates the induction of phase II detoxifying enzyme genes through antioxidant response elements. Biochem Biophys Res Commun 236, 313–322 (1997).

19. Zhang, D. D. Thirty years of NRF2: advances and therapeutic challenges. Nat Rev Drug Discov (2025) doi:10.1038/s41573-025-01145-0.

20. Itoh, K. et al. Keap1 represses nuclear activation of antioxidant responsive elements by Nrf2 through binding to the amino-terminal Neh2 domain. Genes Dev 13, 76–86 (1999).

21. Pfefferlé, M. & Vallelian, F. Transcription Factor NRF2 in Shaping Myeloid Cell Differentiation and Function. Adv Exp Med Biol 1459, 159–195 (2024).

22. Kobayashi, E. H. et al. Nrf2 suppresses macrophage inflammatory response by blocking proinflammatory cytokine transcription. Nat Commun 7, 11624 (2016).

23. Vallelian, F. et al. Heme-stress activated NRF2 skews fate trajectories of bone marrow cells from dendritic cells towards red pulp-like macrophages in hemolytic anemia. Cell Death Differ 29, 1450–1465 (2022).

24. Thimmulappa, R. K. et al. Nrf2 is a critical regulator of the innate immune response and survival during experimental sepsis. J Clin Invest 116, 984–995 (2006).

25. Pfefferlé, M. et al. Hemolysis transforms liver macrophages into antiinflammatory erythrophagocytes. J Clin Invest 130, 5576–5590 (2020).

26. Vallelian, F., Buehler, P. W. & Schaer, D. J. Hemolysis, free hemoglobin toxicity, and scavenger protein therapeutics. Blood 140, 1837–1844 (2022).

27. Consonni, F. M. et al. Heme catabolism by tumor-associated macrophages controls metastasis formation. Nat Immunol 22, 595–606 (2021).

28. Humar, R., Schaer, D. J. & Vallelian, F. Erythrophagocytes in hemolytic anemia, wound healing, and cancer. Trends Mol Med 28, 906–915 (2022).

29. Ingoglia, G. et al. Line-selective macrophage activation with an anti-CD40 antibody drives a hemophagocytic syndrome in mice. Blood Adv 4, 2751–2761 (2020).

30. Buhtoiarov, I. N. et al. CD40 ligation activates murine macrophages via an IFN-gamma-dependent mechanism resulting in tumor cell destruction in vitro. J Immunol 174, 6013–6022 (2005).

31. Bill, R. et al. macrophage polarity identifies a network of cellular programs that control human cancers. Science 381, 515–524 (2023).

32. Liu, Z. et al. Fate Mapping via Ms4a3-Expression History Traces Monocyte-Derived Cells. Cell 178, 1509–1525.e19 (2019).

33. Shah, R., Ibis, B., Kashyap, M. & Boussiotis, V. A. The role of ROS in tumor infiltrating immune cells and cancer immunotherapy. Metabolism 151, 155747 (2024).

34. Rao, J. et al. Macrophage nuclear factor erythroid 2-related factor 2 deficiency promotes innate immune activation by tissue inhibitor of metalloproteinase 3-mediated RhoA/ROCK pathway in the ischemic liver. Hepatology 75, 1429–1445 (2022).

35. Keleku-Lukwete, N. et al. Nrf2 activation in myeloid cells and endothelial cells differentially mitigates sickle cell disease pathology in mice. Blood Adv 3, 1285–1297 (2019).

36. Taguchi, K. et al. Genetic analysis of cytoprotective functions supported by graded expression of Keap1. Mol Cell Biol 30, 3016–3026 (2010).

37. Steele, N. G. et al. Multimodal Mapping of the Tumor and Peripheral Blood Immune Landscape in Human Pancreatic Cancer. Nat Cancer 1, 1097–1112 (2020).

38. Wang, W. et al. Identification of hypoxic macrophages in glioblastoma with therapeutic potential for vasculature normalization. Cancer Cell 42, 815–832.e12 (2024).

39. Chu, X. et al. Integrative single-cell analysis of human colorectal cancer reveals patient stratification with distinct immune evasion mechanisms. Nat Cancer 5, 1409–1426 (2024).

40. Sanin, D. E., et al. A common framework of monocyte-derived macrophage activation. Sci Immunol 7, eabl7482 (2022).

41. Kerzel, T. et al. In vivo macrophage engineering reshapes the tumor microenvironment leading to eradication of liver metastases. Cancer Cell 41, 1892–1910.e10 (2023).

42. Pastushenko, I. & Blanpain, C. EMT Transition States during Tumor Progression and Metastasis. Trends Cell Biol 29, 212–226 (2019).

43. Gross, E. T. E. et al. Immunosurveillance and immunoediting in MMTV-PyMT-induced mammary oncogenesis. Oncoimmunology 6, e1268310 (2017).

44. Beatty, G. L. et al. CD40 agonists alter tumor stroma and show efficacy against pancreatic carcinoma in mice and humans. Science 331, 1612–1616 (2011).

45. Pfefferlé, M. et al. Antibody-induced erythrophagocyte reprogramming of Kupffer cells prevents anti-CD40 cancer immunotherapy-associated liver toxicity. J Immunother Cancer 11, (2023).

46. Park, M. D., Silvin, A., Ginhoux, F. & Merad, M. Macrophages in health and disease. Cell 185, 4259–4279 (2022).

47. Vonderheide, R. H. CD40 Agonist Antibodies in Cancer Immunotherapy. Annu Rev Med 71, 47–58 (2020).

48. Salomon, R. & Dahan, R. Next Generation CD40 Agonistic Antibodies for Cancer Immunotherapy. Front Immunol 13, 940674 (2022).

49. Dangaj, D. et al. Cooperation between Constitutive and Inducible Chemokines Enables T Cell Engraftment and Immune Attack in Solid Tumors. Cancer Cell 35, 885–900.e10 (2019).

50. House, I. G. et al. Macrophage-Derived CXCL9 and CXCL10 Are Required for Antitumor Immune Responses Following Immune Checkpoint Blockade. Clin Cancer Res 26, 487–504 (2020).

51. Klement, J. D. et al. An osteopontin/CD44 immune checkpoint controls CD8+ T cell activation and tumor immune evasion. J Clin Invest 128, 5549–5560 (2018).

52. Hegde, S., et al. Myeloid progenitor dysregulation fuels immunosuppressive macrophages in tumors. bioRxiv (2024) doi:10.1101/2024.06.24.600383.

53. Kilian, M. et al. MHC class II-restricted antigen presentation is required to prevent dysfunction of cytotoxic T cells by blood-borne myeloids in brain tumors. Cancer Cell 41, 235–251.e9 (2023).

54. Hänggi, K. et al. Interleukin-1α release during necrotic-like cell death generates myeloid-driven immunosuppression that restricts anti-tumor immunity. Cancer Cell 42, 2015–2031.e11 (2024).

55. Li, C. et al. Tumor-associated macrophages: potential therapeutic strategies and future prospects in cancer. J Immunother Cancer 9, (2021).

56. van Elsas, M. J. et al. Immunotherapy-activated T cells recruit and skew late-stage activated M1-like macrophages that are critical for therapeutic efficacy. Cancer Cell 42, 1032–1050.e10 (2024).

57. Casazza, A. et al. Impeding macrophage entry into hypoxic tumor areas by Sema3A/Nrp1 signaling blockade inhibits angiogenesis and restores antitumor immunity. Cancer Cell 24, 695–709 (2013).

58. Telarovic, I. et al. Delayed tumor-draining lymph node irradiation preserves the efficacy of combined radiotherapy and immune checkpoint blockade in models of metastatic disease. Nat Commun 15, 5500 (2024).

59. Kong, X. et al. Enhancing Nrf2 pathway by disruption of Keap1 in myeloid leukocytes protects against sepsis. Am. J. Respir. Crit. Care Med. 184, 928–938 (2011).

60. Reddy, N. M., Potteti, H. R., Mariani, T. J., Biswal, S. & Reddy, S. P. Conditional deletion of Nrf2 in airway epithelium exacerbates acute lung injury and impairs the resolution of inflammation. Am. J. Respir. Cell Mol. Biol. 45, 1161–1168 (2011).

61. Pfefferlé, M. et al. Acute Hemolysis and Heme Suppress Anti-CD40 Antibody-Induced Necro-Inflammatory Liver Disease. Front Immunol 12, 680855 (2021).

62. Vallelian, F. et al. Revisiting the putative role of heme as a trigger of inflammation. Pharmacol Res Perspect 6, e00392 (2018).

63. Dobin, A. et al. STAR: ultrafast universal RNA-seq aligner. Bioinformatics 29, 15–21 (2013).

64. Danecek, P. et al. Twelve years of SAMtools and BCFtools. Gigascience 10, giab008 (2021).

65. Liao, Y., Smyth, G. K. & Shi, W. The R package Rsubread is easier, faster, cheaper and better for alignment and quantification of RNA sequencing reads. Nucleic Acids Res. 47, e47 (2019).

66. Love, M. I., Huber, W. & Anders, S. Moderated estimation of fold change and dispersion for RNA-seq data with DESeq2. Genome Biol. 15, 550 (2014).

67. Lun, A. T. L., McCarthy, D. J. & Marioni, J. C. A step-by-step workflow for low-level analysis of single-cell RNA-seq data with Bioconductor. F1000Res. 5, 2122 (2016).

68. Korsunsky, I. et al. Fast, sensitive and accurate integration of single-cell data with Harmony. Nat. Methods 16, 1289–1296 (2019).

69. Hie, B., Bryson, B. & Berger, B. Efficient integration of heterogeneous single-cell transcriptomes using Scanorama. Nat Biotechnol 37, 685–691 (2019).

70. Badia-I-Mompel, P., et al. decoupleR: ensemble of computational methods to infer biological activities from omics data. Bioinform Adv 2, vbac016 (2022).

71. Fang, Z., Liu, X. & Peltz, G. GSEApy: a comprehensive package for performing gene set enrichment analysis in Python. Bioinformatics 39, btac757 (2023).

72. Wolf, F. A., Angerer, P. & Theis, F. J. SCANPY: large-scale single-cell gene expression data analysis. Genome Biol. 19, 15 (2018).

